# Reconstruction of hundreds of reference ancestral genomes across the eukaryotic kingdom

**DOI:** 10.1101/2022.02.17.480882

**Authors:** Matthieu Muffato, Alexandra Louis, Nga Thi Thuy Nguyen, Joseph Lucas, Camille Berthelot, Hugues Roest Crollius

## Abstract

Ancestral sequence reconstruction is a fundamental aspect of molecular evolution studies, and can trace small-scale sequence modifications through the evolution of genomes and species. In contrast, fine-grained reconstructions of ancestral genome organisations are still in their infancy, limiting our ability to draw comprehensive views of genome and karyotype evolution. Here we reconstruct the detailed gene contents and organisations of 624 ancestral vertebrate, plant, fungi, metazoan and protist genomes, 183 of which are near-complete chromosomal reconstructions. Reconstructed ancestral genomes are similar to their descendants in terms of gene content as expected and agree precisely with reference cytogenetic and *in silico* reconstructions when available. By comparing successive ancestral genomes along the phylogenetic tree, we estimate the intra- and inter-chromosomal rearrangement history of all major vertebrate clades at high resolution. This freely available resource introduces the possibility to follow evolutionary processes at genomic scales in chronological order, across multiple clades and without relying on a single extant species as reference.

## Introduction

Biological sequences have long been recognized as a document of evolutionary history^1^, where accumulated mutations record relationships between species as well as the dynamics underlying their evolution. Given sufficient genetic information across species, the temporal accumulation of these mutations can be traced back in time to reconstruct sequences and genomes in their long-lost common ancestors. These ancestral reconstructions are the backbone of much of today’s methodologies in molecular evolution, including phylogenetic trees^2–4^ and sequence selection tests^5,6^. The reconstruction of ancestral sequences, and especially genes, has been extensively studied since the dawn of sequencing: mature methods exist to retrace the history of sequence substitutions and leverage changes in substitution dynamics to answer specific evolutionary questions. However, DNA mutations are not limited to sequence substitutions: genomes are also affected by larger scale mutational events such as duplications, deletions, sequence inversions or chromosomal rearrangements, all of which can affect genome function, species fitness, and evolution. In extant species, large-scale mutations are a major determinant of disease as they can disrupt functional sequences^7–9^ as well as reorganize functional structures within the genome^10–12^. From an evolutionary viewpoint, large-scale mutations are a well-documented source of innovations: they can produce new genetic combinations that contribute phenotypic novelty^13,14^, but can also have more indirect effects such as locally suppressing recombination^15,16^, favouring allele hitchhiking and rapid selection^17,18^. For example, genomic rearrangements have been shown to associate with changes in brain gene expression between humans and chimpanzees^19^, to underlie the evolution of intersexual development in moles^20^ and variations in reproductive morphs in ruffs^21^. Despite their tremendous functional and evolutionary importance, large scale mutational events are less extensively studied and not as well-understood than sequence substitutions. In particular, the reconstruction of ancestral genomes and karyotypes lags behind that of ancestral sequences, making it difficult to study the evolutionary dynamics and impact of rearrangements, duplications and deletions over many species and within rigorous theoretical frameworks.

With the advent of massive sequencing projects ambitioning to obtain high-quality reference genomes for thousands of species across all kingdoms of life^22^, evolutionary genomics faces both fresh opportunities and serious challenges to integrate this flow of data into usable comparative frameworks. Along with whole-genome alignments^23^, ancestral genome and karyotype reconstructions across large clades is one of the most promising outcomes of these projects. The goal of these reconstructions is to provide a plausible organization of genomic sequences in one or many extinct common ancestors of a group of species of interest. Several paleogenomics strategies have been explored to reconstruct the sequence content and ordering of ancestral genomes. Methods based on double-cut-and-join (DCJ) algorithms endeavour to reconstruct rearrangement scenarios resulting in observed extant genome structures^24,25^. These methodologies become increasingly computationally expensive and in many cases intractable for sets of large, complex genomes, which at this time has only been overcome by reducing significantly reconstruction resolution^26–28^. Other methods attempt to reconstruct a parsimonious sequence ordering in the ancestor based on orthologous sequence adjacencies in extant genomes, under the assumption that genomic rearrangements are unlikely to result in the same sequence organization several times independently. These methods can be applied to different types of markers, typically either alignable sequence blocks or individual genes, and are appropriate for small^29^ and large genomes such as vertebrates or plants^30,31^. However, it remains unclear whether current methods can provide high-resolution reconstructions and scale to the large genomic resources available in comparative genomics databases. At this time, only two ancestral genomic reconstruction resources are widely available to the community: AncestralGenomes^32^, which provides 111 ancestral gene content reconstructions, but not their order (‘bags of genes’), and DESCHRAMBLER^33^, which offers chromosome-complete reconstructions for 7 mammal and 14 bird ancestors, but with limited sub-chromosomal resolution (100-300 kb sequence blocks) and dependent on a reference genome. Here, we introduce a new resource containing 624 ancestral genomes reconstructed over the vertebrate, plant, fungi, metazoan and protist clades, at gene-scale resolution, where a third of the ancestral genomes reach chromosomal-complete assemblies. This drastic change in magnitude is powered by an iterative, parsimony-based ancestral genome reconstruction algorithm, named AGORA, which we describe here. We show that AGORA is efficient, flexible and scales to integrate hundreds of large genomes and to reconstruct their common ancestors at every node in the species phylogeny with relatively modest computational costs. Along with the open-source algorithm, all precomputed ancestral genome reconstructions are publicly available within the Genomicus^34,35^ database (https://www.genomicus.bio.ens.psl.eu/genomicus), and benefit from the full browsing and comparative genomics tool infrastructure of the database. The database is regularly updated since 2010 to reflect reference genome improvements and represents a perennial resource for high-quality, high-resolution ancestral genomes for the molecular evolution community across disciplines and model phylogenetic clades.

## Results

### A vast resource of eukaryotic reference ancestral genomes for evolutionary genomics

To facilitate the investigation of chromosomal and local genome dynamics across evolution, we have developed an extensive resource of ancestral genome reconstructions that spans large portions of the eukaryotic tree of life. This resource is based on an algorithm named AGORA (Algorithm for Gene Order Reconstruction in Ancestors), which computes highly contiguous, near-exhaustive reconstructions of the ancestral gene order at every bifurcation in the species tree, based on gene order information in the extant species of the clade (Figure 1a). While AGORA can be installed as a standalone package for tailored research applications, we routinely precompute and release the complete set of ancestral vertebrate genomes for every update of the Ensembl database as well as for a broad selection of plant and fungi clades as part of the Genomicus synteny database^36^. At the time of submission, Genomicus contains a total of 624 ancestral genomes readily available for download across the Vertebrates, Plants, Metazoa, Protists and Fungi databases (Supplementary table S1). These ancestral genomes can be explored and manipulated using the different utilities of the Genomicus web server^36^ to perform karyotype comparisons, extraction and evolutionary tracing of conserved synteny blocks (Figure 1b), and local gene-to-gene synteny visualization across ancestral and extant species (Figure 1c).

**Figure 1.**
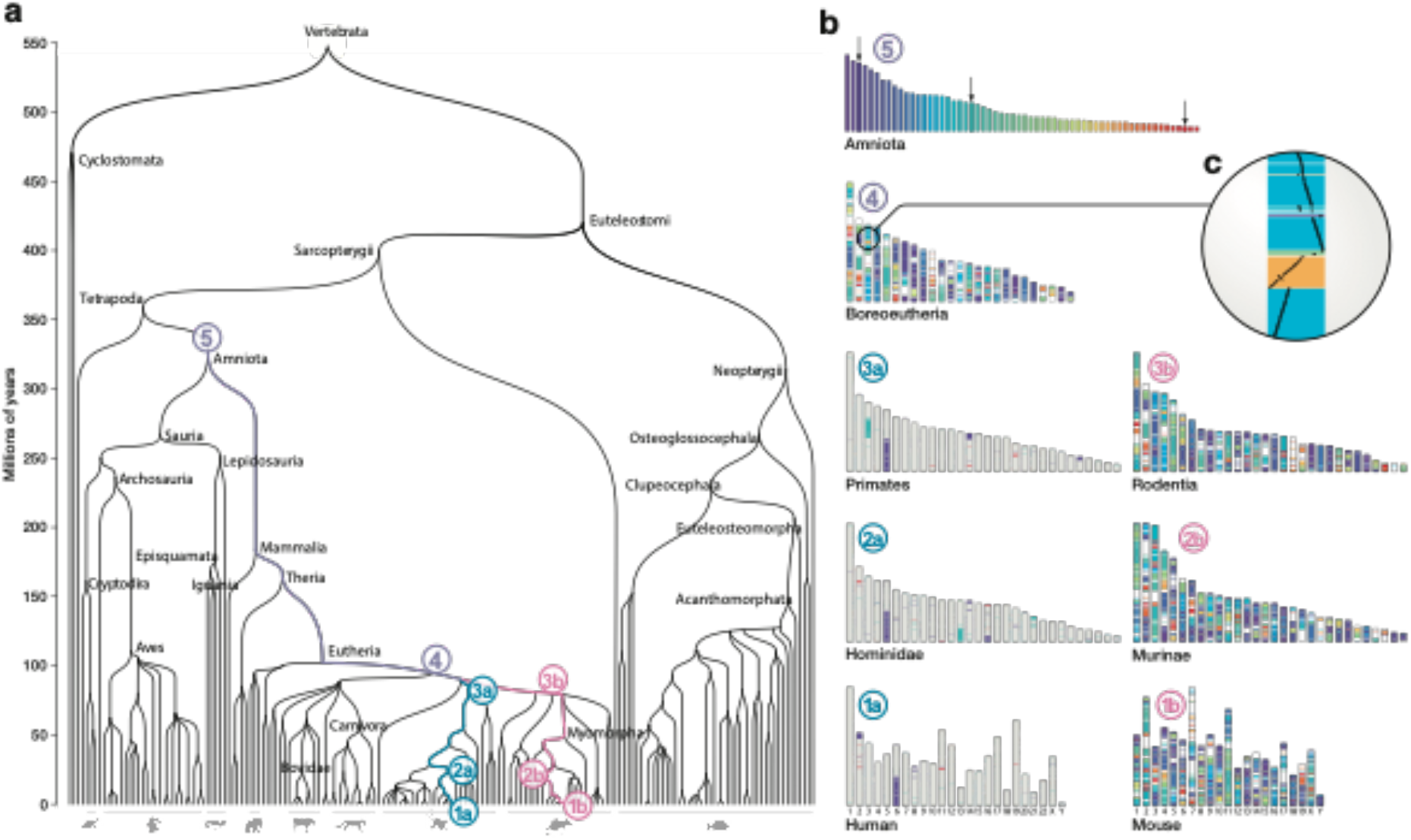
Reconstructing vertebrate ancestral genomes. **a.** Species phylogeny of vertebrates encompassing genomes stored in Ensembl version 92 with indications of the eight ancestral genomes detailed in panels B and the evolutionary path that they mark out. **b.** High resolution ideograms of ancestral genome reconstructions starting from the *Amniota* genome (5) and the descendant *Boreoeutheria* genome (4), where a region on the 3^rd^ chromosome is expanded to highlight the evolution of gene organisation with respect to the *Amniota* genome. In the primate lineage (3a, 2a, 1a) only the evolution of the three *Amniota* chromosomes indicated by an arrow are depicted in colour, while in the *Rodentia* lineage (3b, 2b, 1b) the evolution of all *Amniota* chromosomes is shown.

### AGORA: a fast and efficient algorithm to reconstruct ancestral gene order at every node in a phylogenetic tree

AGORA is a parsimony-based algorithm that estimates the content and order of genes in the ancestor of a group of extant species for which reference genomes are available (Figure 2, Figure S1). Briefly, the method iteratively extracts commonalities between pairs of extant genomes to infer characteristics inherited from their last common ancestor, and present in every ancestor along the evolutionary branches leading to each extant genome. AGORA takes as input a forest of gene phylogenetic trees, corresponding to all the gene families present in the extant genomes with their orthology and paralogy relationships, and the gene orders in each extant genome. First, AGORA uses the phylogenies of extant genes to infer the gene content of every ancestor along the species tree (Figure S2). Second, AGORA compares the gene orders of every pair of extant species to identify orthologous genes adjacent and in the same orientation in both species, and presumably inherited from their last common ancestor (Figure 2a). For every ancestor in the species tree, the algorithm extracts the subset of informative pairwise extant species comparisons (Figure 2b) and integrates the gene adjacency comparisons into a weighted graph, where nodes represent ancestral genes, and edges adjacencies supported by pairwise extant species comparisons. The weights correspond to the number of comparisons supporting that these genes were adjacent in this ancestor (Figure 2c,d). Ideally, this process would result in a linear graph representing the ancestral gene order, as genome rearrangements are unlikely to produce the same gene adjacencies independently in different lineages^37–40^. However, errors in the resolution of orthologs and paralogs in the original gene trees can result in branching in the graph. AGORA linearizes the graph by iteratively removing low-weight edges to obtain a parsimonious reconstruction of the oriented gene order in the ancestral genome (Figure 2e). AGORA includes extensions of this algorithm to deal with larger errors in the input gene trees, by identifying a set of constrained genes that are close to being single copy in most species, and can be reliably used for gene order reconstruction. In this mode, AGORA adds the non-constrained genes in a second stage. The algorithm is presented in detail in the Supplementary Information (Figures S1-9). The *in-silico* performances of AGORA have been tested on a previously-used benchmark of genome evolution simulations^33^, achieving 98.9% agreement with the reference (sensitivity 99.3%; precision 99.6%; Methods), similar to other state-of-the-art ancestral genome reconstruction methods^33^. On a different, more realistic, benchmark based on simulations that are not restricted to single-copy genes, AGORA achieves 95.4% agreement, while DESCHRAMBLER’s performance drops to 68.6% (Supplementary Material), highlighting AGORA’s ability to successfully deal with gene duplications and other complex evolutionary scenarios.

**Figure 2.**
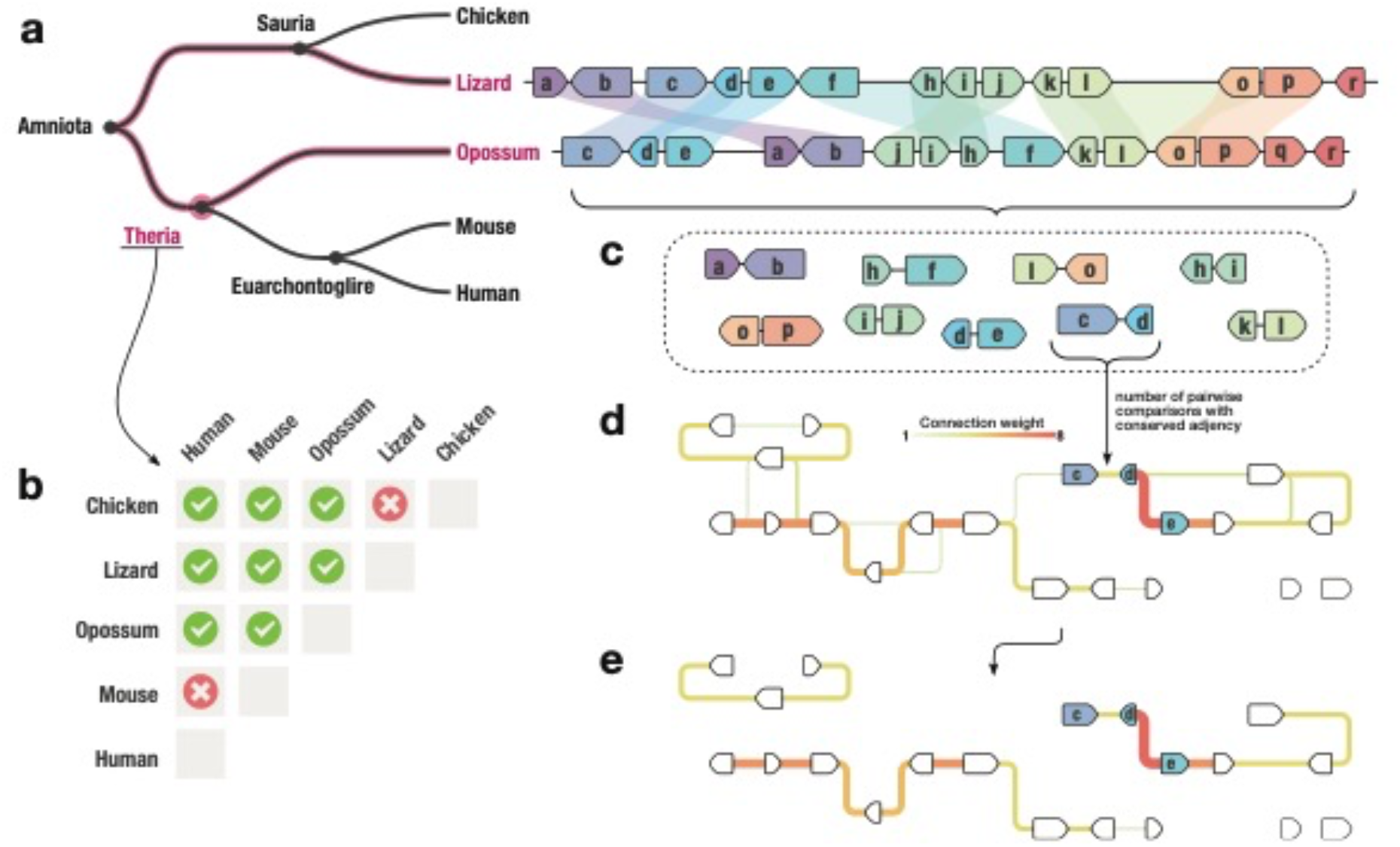
Principle of the AGORA approach. **a**. Conserved gene adjacencies are identified between all genome pairs that are informative for a given target ancestral genome. Here a portion of the Lizard and Opossum genomes are shown, with gene adjacencies joined by a pale coloured shape when conserved, and thus supporting their prior occurrence in Theria. **b.** All comparisons between genomes that are joined in an evolutionary path intersecting the target ancestor are informative (green ticks) while comparisons between genomes that diverged after the target ancestor are uninformative (red crosses). **c-d**. Conserved adjacencies observed in each pairwise comparison **(c)** are collected in a graph structure **(d)** where nodes are genes and links are conserved adjacencies weighted by the number of times they have been observed in pairwise genome comparisons. **e**. The linearisation of this graph by traversing the links of maximal weight provides contiguous and parsimonious ancestral gene order reconstructions.

In practice, AGORA is highly flexible as it only requires the protein-coding gene annotations of the extant species and the set of precomputed gene trees in a standard format, which can be downloaded from a variety of genome resource initiatives for many species groups. AGORA can be used with other markers than protein-coding genes, such as conserved non-coding elements; however, due to unreliability of phylogenetic trees for those sequences, we recommend limiting the reconstructions to the order of protein-coding genes for best performances. AGORA can also be used iteratively to assemble blocks of markers and scaffold them over several rounds of reconstruction into larger contiguous ancestral regions (CARs). We propose several workflows customized for different clades and applications as part of the AGORA package (Figure S1).

Here, to demonstrate the capabilities of AGORA, we use two datasets from distant eukaryotic clades, with different numbers of species, genes, and variable gene tree reliability: i) a dataset of 93 vertebrates and 5 outgroups and their 23,528 gene trees, including a total of 1,814,614 extant protein-coding genes, and leading to the reconstruction of 81 ancestral genomes; and ii) a dataset of 58 plant genomes and 8 outgroups, corresponding to 48 ancestral genomes (Methods; Supplementary table S4 and Supplementary File 1).

### Reconstructions of highly contiguous, chromosome-scale assemblies of key ancestral genomes

For every ancestral genome, we provide two valuable results: the gene set, and an assembly of their ancestral organization. To evaluate the completeness and accuracy of the ancestral gene sets, we first compare the total number of genes inferred in an ancestor to those of its descendant extant genomes. While very distant genomes can contain widely different numbers of genes, AGORA is designed to be used within clades where synteny is reasonably conserved, such as vertebrates, grasses or *Saccharomycetales* yeasts, and where genomes typically contain similar numbers of genes. We find that our methodology accurately estimates ancestral gene contents that are consistent with those of the descending clades, up to evolutionary distances of over 300 My (Figure 3a). We also find that the vast majority of clade-relevant BUSCO (Benchmark Universal Single Copy Orthologs)^41^ reference sets are present as single copy genes in our inferred ancestral gene sets (Figure 3b). In addition, we also confronted our inferred ancestral gene contents for seven key vertebrate ancestors to those calculated by Ancestral Genomes, another effort to estimate the ancestral gene content - but not gene order - at different evolutionary nodes^29^. Ancestral Genomes relies on the PANTHER database^42^ and therefore uses an independent set of extant genomes and gene trees. AGORA and Ancestral Genomes both infer highly similar gene contents for the same ancestors (Figure 3c).

**Figure 3.**
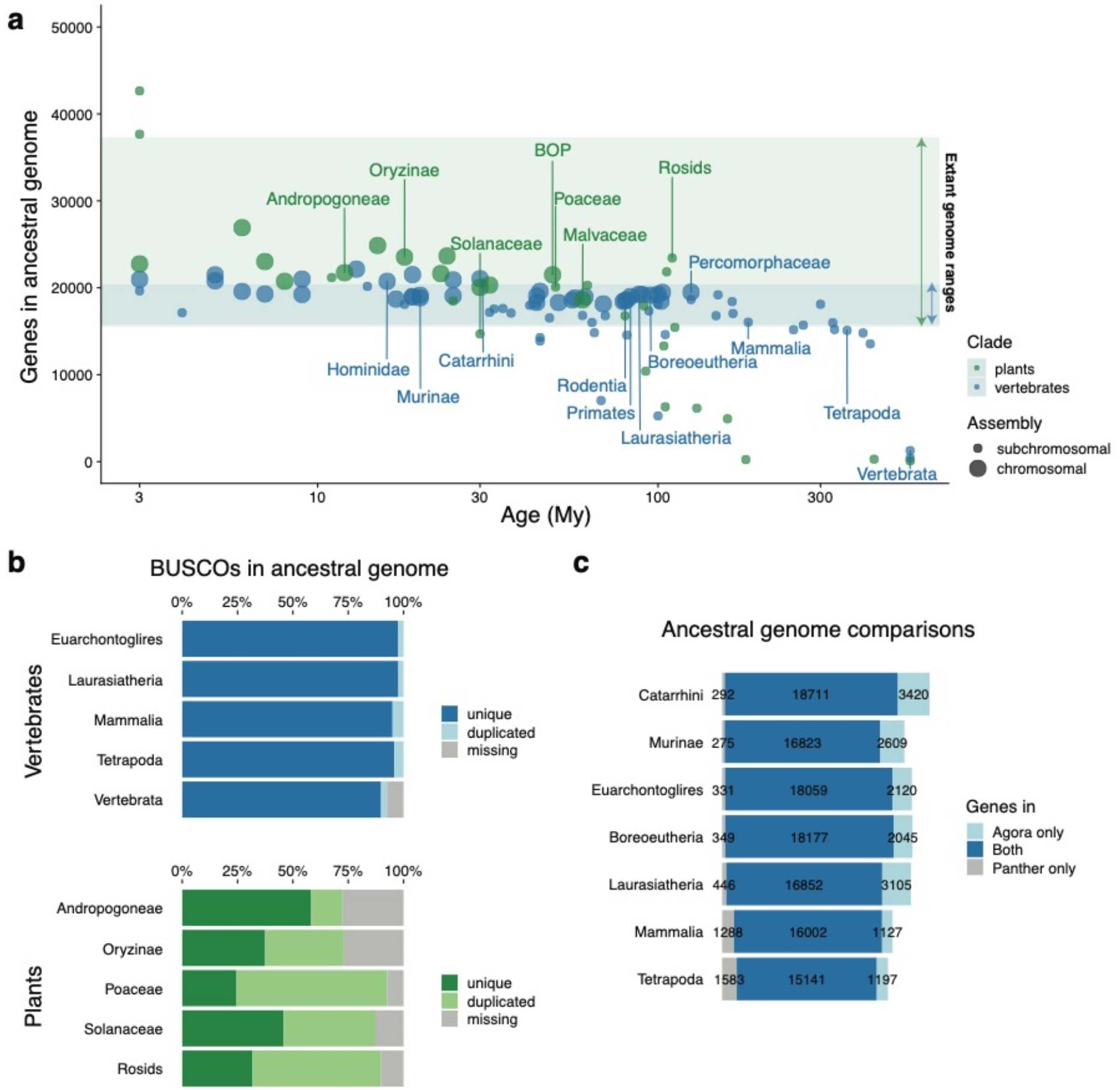
Completion of ancestral genomes reconstructed by AGORA. **a**. Gene content and assembly continuity of 77 vertebrate and 33 plant ancestral genomes reconstructed by AGORA. The ranges of gene contents of extant vertebrate and plant genomes are highlighted as blue and green shading, respectively. Very young (< 2 My) or very old (> 550 My) ancestors are not represented. Chromosomal and subchromosomal assemblies are as defined in Methods. **b.** Representation of BUSCO genes in AGORA’s ancestral genomes. Plant genomes, which have undergone rounds of whole-genome duplications, frequently contain a large fraction of duplicated genes. **c.** Comparison of ancestral gene contents inferred by reconstructions from AGORA and PANTHER.

The other output of AGORA is the reconstruction of the putative gene order in each ancestral genome along the species tree. The quality of an ancestral genome reconstruction can be evaluated by two criteria: contiguity, and consistency with evolutionary and biological evidence. Contiguity represents the size of the genomic regions that can be assembled into contiguous ancestral regions (CARs), akin to measures of assembly quality for reference genome sequences. For 37 vertebrate ancestral genomes and 13 plant ancestral genomes in our test set, we obtain chromosome-scale assemblies with a small number of long CARs containing hundreds of ordered and oriented genes, corresponding to a best approximation of the ancestral karyotype (Figure 3a). These chromosome-level assemblies include over 70% of the ancestral genes, which is comparable to well-assembled extant reference genomes in those clades (Figure S10). Most other ancestral genomes are assembled into fewer than 100 subchromosomal gene blocks containing over 70% of the ancestral gene content (Figure S11).

As expected, the contiguity of ancestral genome reconstructions is overall high in recent ancestors and decreases sharply after 100 My, decaying to large numbers of short, unassembled gene blocks for very ancient ancestors such as the *Tetrapoda* and *Vertebrata* ancestors (Figure 3a). However, perhaps counterintuitively, AGORA performs better in some key older ancestors than in comparatively younger ancestral genomes. For example, the genome of *Boreoeutheria*, the ancestor of most placental mammals (approx. 95 Mya), is a near-complete assembly comprising 25 large CARs covering 18,430 genes (80% of the total ancestral genome), while the genome of *Afrotheria*, the ancestor of elephant and hyrax (approx. 90 Mya), is appreciably less contiguous with 70% of genes in 83 CARs. This reflects the position of these ancestors in the species tree relative to the sampling of sequenced extant genomes. As previously demonstrated^43^, ancestors that precede evolutionary radiations are ideally positioned for ancestral genome reconstruction, as their many outgroup and descendant lineages offer a large number of informative pairwise comparisons (N_i_). Overall, AGORA’s ancestral reconstruction contiguity correlates with the N_i_/age ratio (Figure S12). Because sequencing efforts have largely targeted organisms within species-rich phyla such as placental mammals or monocotyledon plants, the key ancestors to these widely-studied subclades are particularly well-reconstructed by our methodology, which will be of high value to evolutionary and functional studies. Ultimately however, with the advent of massive sequencing undertakings such as the Vertebrate Genome Project, genome documentation in under-sampled clades will increase dramatically and we expect that most ancestral genomes in the Genomicus database will eventually become chromosome-level assemblies.

### Recapitulation of evolutionary evidence and curated *in-silico* paleogenomics reconstructions

Accuracy of ancestral genome reconstructions is appreciably more difficult to evaluate than completion, as the true ancestral genome sequences are inaccessible at the evolutionary scales we study. However, several ancestral genomes have garnered longstanding interest from the evolutionary genomics community, resulting in a large body of biological evidence regarding their overall organization. In vertebrates, one of the most studied ancestral genomes is *Boreoeutheria*, the 95 My-old ancestor to most placental mammals including primates, rodents, hooved mammals and carnivores, with the exception of afrotherians (elephants) and xenarthrans (sloths, anteaters, armadillos), along with the *Eutheria* ancestor (102 My old, ancestral to boreoeutherian mammals and afrotherians) and the *Simian* ancestor (45 My old, ancestral to platyrrhine and catarrhine primates). Landmark ancestral *Eutheria, Boreoeutheria* and *Simian* karyotypes have previously been reconstructed by integrating dozens of mammalian homology comparisons using fluorescent DNA probes, a technique known as chromosome painting^44,45^. This analysis suggested that the ancestral placental genome comprised 23 pairs of chromosomes, and traced the large-scale rearrangements that resulted into the karyotypic arrangement of the human genome. The *Boreoeutheria* ancestral genome organisation inferred by AGORA contains 25 large CARs and is highly congruent with the cytogenetics-based reference karyotype (Figure 4a). AGORA recovers all ancestral chromosomal arrangements supported by cytogenetics evidence without requiring manual assembly or curation. The only exception is the ancestral linkage of human chromosomes 10 and 12 alleged by cytogenetics data (Figure 4b-d), which is supported neither by AGORA, nor by the state-of-the-art reconstruction by DESCHRAMBLER or other *in-silico* ancestral genome reconstruction methods^33^. Detailed manual investigation of inconsistencies between the ancestral reconstructions by AGORA and the cytogenetics references reveals that most differences are the result of the lower resolution of the chromosomal painting methodology, and confirm our proposed assembly (Figures S14-15). At the infrachromosomal scale, we found that the genomic organization of the *Boreoeutheria* genome inferred by AGORA is in near perfect agreement with that of DESCHRAMBLER (Figure 4c; Methods). However, our reconstructed *Boreoeutheria* genome is more complete and includes the ancestral locations of an additional 2,023 genes (8% of the ancestral gene set). Altogether, these results support that the gene-based reconstruction algorithm of AGORA is highly consistent with current ancestral reconstruction methods, while providing a significant increase in resolution for the study of local genomic events.

**Figure 4.**
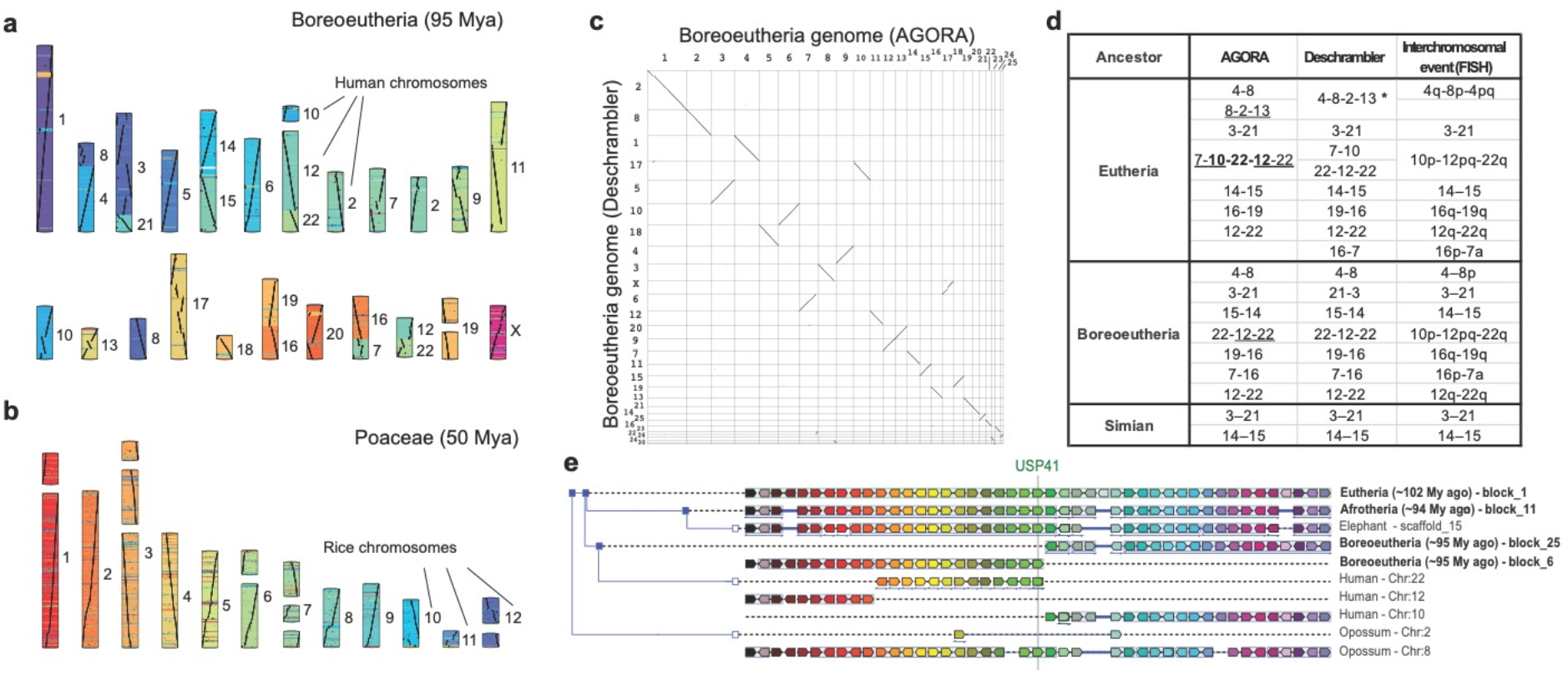
AGORA ancestral genome reconstructions compared to extant genomes and state-of-the-art ancestral reconstructions. **a**. The *Boreoeutheria* karyotype inferred by AGORA (25 largest CARs), coloured according to gene locations on human chromosomes, as indicated to the right of each CAR. **b**. The *Poaceae* karyotype inferred by AGORA (19 largest CARs), coloured according to gene locations on *Oryza sativa* chromosomes, as indicated to the right of each CAR. **c**. Collinearity of the *Boreoeutheria* ancestral genomes reconstructed by AGORA and DESCHRAMBLER. **d.** Comparisons of computational reconstructions by AGORA and DESCHRAMBLER, and Zoo-FISH linkage groups inferred for three key mammalian ancestors. Human chromosomes in ancestral linkage are indicated with hyphens. The *Eutheria* bolded linkage group 10-22-12 is documented in more detail in **(e)**. Underlined linkage groups are documented in Fig S14. DESCHRAMBLER reconstructs a linkage group between parts of human chromosomes 4, 8, 12 and 3 (asterisk) in disagreement with FISH evidence and AGORA when used on Ensembl v.92 data; however, this linkage group is also reconstructed by AGORA on Ensembl data v.102, suggesting an ambiguous ancestral linkage state (Fig S15). **e**. Gene adjacencies around USP41 in extant species support that fragments of human chromosomes 10, 22 and 12 were linked in the *Eutheria* ancestor. Ortholog genes are shown in matching colours. Opossums and elephants have both retained the ancestral organisation at this locus, which has been rearranged in the human genome.

We also compared the genome of *Poaceae*, the 50 My-old ancestor of grasses, reconstructed by AGORA to an earlier reference ancestral karyotype^46^ obtained by another parsimony-based method to reconstruct ancestral adjacencies^30^. Again, the ancestral genome reconstructed by AGORA closely recapitulates the state-of-the-art knowledge regarding the organization of the ancestral grass karyotype (Figure 4e), while providing access to a fine-scale reconstruction of the ancestral gene order that was unavailable to date.

### A scalable framework to integrate genomic data across entire phylogenies

A major strength of AGORA resides in its ability to compute the gene order of every ancestor in a phylogeny using different subsets of the same extant genome comparisons. In a context where new species genomes are being sequenced with increasing speed and accuracy, comparative genomics need methods that can integrate evolutionary information along the species tree and across lineages without relying on a single extant genome as reference. Using the legacy architecture of the Genomicus synteny database^34,35^, which is updated with every new release of the Ensembl database, we tested how our methodology scales with the number of extant reference genomes available as well as their quality (Supplementary figure S13). Ensembl Compara v.101 included the reference sequences of 264 vertebrate species and 5 outgroups, for a total of 5,539,325 extant protein-coding genes organised into 62,478 gene trees. Using this information, AGORA reconstructs a total of 265 ancestral genomes along the species tree in 6 hour and 50 minutes on a Linux machine with 4 CPUs and ~80 GB RAM (Supplementary table S2). AGORA is therefore computationally inexpensive and can be run on a desktop machine for small to medium datasets. However, AGORA can also be parallelized and is optimized for usage on a computing cluster for large applications and database updates (Supplementary Information).

Overall, the quality of key ancestral genomes increases as new extant genomes are included in the database (Supplementary figure S13). The introduction of high-quality reference genomes in under-represented clades over time has contributed to the reconstruction of previously inaccessible ancestors, such as *Strepsirrhini*, the ancestor of lemurs, bushbabies and lorises, and more recently *Chiroptera*, the ancestor of bats. Interestingly, we observed that even the inclusion of low-coverage, fragmented reference genomes significantly improves ancestral genome reconstructions. This is likely because different reference genomes are generally assembled independently, and assembly errors rarely produce the same erroneous gene arrangements from one genome to the next. As AGORA only considers conserved gene adjacencies as potentially ancestral, the additional information from correctly assembled scaffolds offsets the noise introduced by assembly errors, which are discarded as not conserved. We therefore argue that the inclusion of low-cost, fragmented reference genomes in comparative genomics databases serves a purpose beyond gene-based analyses.

### Ancestral genomes as backbones for evolutionary studies

In this section, we illustrate how our inferred ancestral genomes can serve a powerful resource for genome evolution studies. As a case study, we used ancestral reconstructions to investigate the patterns of karyotypic rearrangements that occurred during the evolution of mammals, birds and ray-finned fish (Figure 5). These three groups represent the main jawed vertebrate (*Euteleostomi*) lineages, whose respective chromosomal dynamics have been documented using comparative genetics, cytogenetics, and genomics approaches across different taxonomic groups. We selected 73 well-reconstructed ancestors and their 74 extant descendants (15 birds and reptiles, 41 mammals, 18 fish; Methods) from Genomicus Vertebrates database v.102, which contains a total of 269 extant and 265 ancestral genomes. We then compared consecutive genomes on all 131 branches of the phylogenetic tree, representing a combined time of about 5 billion years of independent evolution, and traced gene adjacencies that were rearranged on each branch (Methods). In total, we identified 7,145 rearrangement breakpoints that occurred along the 131 branches (average rate 1.45 breakpoint/million years), most of which are intrachromosomal. We also identified 1,370 interchromosomal rearrangements (translocations, fusions or fissions) with an average rate of 0.28 rearrangement per million year (Figure 5a; Supplementary table S3). Comparing rates per million years, and restricting the analysis to the 105 branches longer than 5 My to avoid small sample distortions, we confirm that birds and reptiles have more stable chromosomal structures than mammals, as previously reported^47,48^, with lower rates of interchromosomal rearrangements (p-value = 3.8e-06, Wilcoxon test; Figure 5b). Fish in turn display higher intrachromosomal breakpoint rates than mammals, birds and reptiles (teleosts versus saurians, p-value = 0.0181; teleosts versus mammals, p-value = 0.0532, Wilcoxon test), consistent with the rediploidisation process following the whole genome duplication that occurred in this phylum^49^, yet they display a uniformly high karyotypic stability.

**Figure 5.**
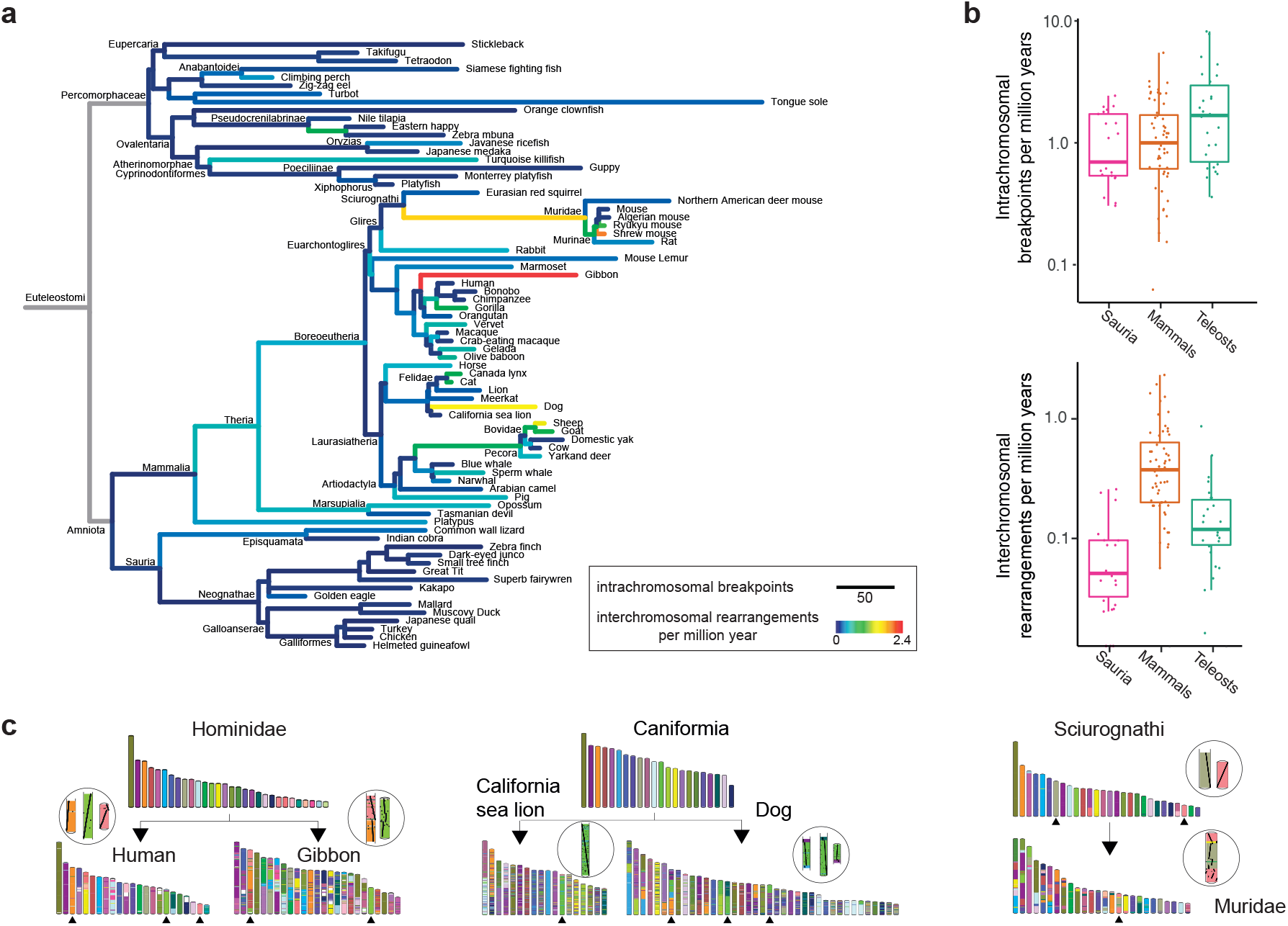
Vertebrate genome evolutionary dynamics. **a.** Phylogenetic tree of 74 extant and 73 ancestral genomes, where branch lengths represent the number of breakpoints computed between successive nodes. Colours represent the rate of interchromosomal rearrangements. Branches in grey connect ancestral genomes their *Euteleostomi* root, which is too fragmented as a genome reconstruction to serve as reference for computing breakpoints and rearrangements. **b**. Distributions of breakpoints and rearrangement rates represented in **(a)**, broken down into three taxonomic groups: saurians (birds and reptiles), mammals and teleosts (fish). **c**. Examples of rearrangements in lineages with notable rates of evolution. Black upward arrow heads point to chromosomes that are shown enlarged in the circles with individual orthologous genes drawn as black dots.

Interestingly, a few branches in placental mammals stand out as having strikingly high rearrangement rates. For instance, the gibbon lineage is the outlier of our analysis, having experienced 60 interchromosomal rearrangements in 25 My, confirming previous observations that this is a fast-evolving lineage compared for example to the human lineage^50^ (Figure 5c). The dog genome was also subject to high rates of rearrangement, especially compared to its sister branch leading to the slowly evolving sea lion genome, which only changed through three chromosome fusions compared to their *Caniformia* ancestor^51^. The lineage leading to *Muridae* is notable for a high rate of intrachromosomal breakpoints combined with multiple interchromosomal rearrangements but associated to a stable chromosome number, consistent with cytogenetic studies of different murid clades^52,53^.

## Discussion

Biology is a historical science, but this historical dimension is often ignored because the records required to document ancestral states are missing. Without this chronological perspective, the reasons why contemporary biological systems are organised as they are will continue to elude us. In practice, this information gap hinders our ability to integrate conclusions across different living models, and to draw the full benefits of comparative genomics. Ancestral genomes are fundamental blocks of the conceptual framework aiming to address this problem. They complement fossils as biological timepoints, as they are a theoretical representation of the precise divergence between two lineages, while fossils represent true extinct species but whose exact phylogenetic position is often unclear. As ancestral genomes encapsulate all the genes present in the ancestral organism as well as their structural organisation, they will enable detailed investigations of developmental and metabolic pathways evolution, such as the expansion and contraction of specific gene families over time; the contribution of genome structure changes to evolutionary transitions and speciations; and the tracing of evolutionary innovations through reorganisation of functional gene arrangements. Additionally, ancestral genomes can act as unique reference points to compare multiple descendant genomes, removing the bias of relying on an extant genome as central reference. This property makes them powerful tools to identify, measure and study lineage-specific genomic events as well as clade-wide trends.

As genome sequencing costs continue to decrease, reference genomes are becoming widely available for model and non-model species alike. At the time of writing, the NCBI database accounts for a total of 8,505 eukaryote, 32,172 bacterial and 1,909 archaeal whole genome sequencing projects, and dedicated efforts such as the Vertebrate Genome Project^54^ promise to deliver extensive phylogenetic coverage across many clades. Integrating sequence and genome organization evolution over such massive phylogenetic samplings remains a challenge. Many phylogenomics projects still rely on sequence alignments as a means to study how genome organization evolves^33,48^. Aligning whole genomes is computationally expensive, and while new methodologies are emerging to step up to the challenge^23,55^, the requirements to handle hundreds of genomes remain out of reasonable reach for many. Additionally, identifying conserved and rearranged regions from whole-genome alignments becomes technically difficult as phylogenetic distance increases, especially in large genomes where a significant fraction of the sequence is non-coding and repetitive. Due to these limitations, the evolution of genome organization is typically studied at large scale, but low resolution, and/or in a limited sampling of species - often those included in publicly available, reference multispecies alignments. Marker-based ancestral genome reconstructions provide an alternative to methods based on whole-genome alignments by relying on gene phylogenies instead, which require much more modest computational infrastructures and scale up to hundreds of genomes with relative ease. As algorithms mature, ancestral genomes such as presented here could become enriched with many more features, including non-coding sequences such as ancestral repeat elements, non-coding RNA genes or regulatory elements, and serve as organizational maps for reconstructed or fossil nucleotide sequences. In the future, as polymorphism information becomes available for more extant species, we may even expect to see ancestral genomes move on from unique references to compendiums, representing structural genomic variation present at any given point in time – opening the door from increasingly sophisticated population genomics models of molecular evolution.

## Methods

### Data collection

Genes and gene trees were downloaded from in Ensembl v.92^56^, and Ensembl Plants v.41^57^. Ensembl v.92 gene trees were edited for poorly supported duplication nodes as previously described^58^, as part of the standard build procedure for the Genomicus synteny database. Of note, this step only marginally improves ancestral genome reconstructions and is not a prerequisite to use AGORA. The species trees for the extant and ancestral genomes from Ensembl v.92, and Ensembl Plants v.41 are described in Supplementary File 1.

### Ancestral genome reconstructions

Ancestral gene sets and gene orders were reconstructed for 82 ancestors on Ensembl v.92 data using AGORA with two passes and a tree parameter of 0.35, and for 41 plant ancestors in two multi-integration passes without tree selection (Supplementary Table S4). The details of the AGORA algorithm, validations by evolutionary simulations, and suggested procedure to select an optimal tree parameter are detailed in Supplementary Information.

### Statistics on ancestral genomes

Ancestral genome contiguity was measured using L70 and G50 metrics. L70 is the smallest number of CARs adding up to 70% of the total genome length, measured in gene units. G50 is the length of the ancestral CAR such that 50% of the total genome length, measured in gene units, is contained in larger CARs. Vertebrate chromosomal assemblies have a L70 < 100 and G50 > 450 and plant chromosomal assemblies have a L70 < 20 and a G50 > 450. These values correspond to well-assembled extant genomes (Figure S10) from these respective clades. Other assemblies were considered subchromosomal.

### Comparisons to reference ancestral gene sets

We downloaded the Benchmark Universal Single Copy Orthologs (BUSCO) sets v.3^41^, based on OrthoDB v.9^59^. BUSCO gene identifiers were converted to Ensembl gene IDs using the conversion tables provided by OrthoDB. A BUSCO orthogroup is a set of near-1-to-1 orthologous genes across sequenced genomes of a relevant phylum. An ancestral gene inferred by AGORA was identified as a BUSCO if two or more of its extant descendant genes are contained in the same orthogroup. When a single ancestral gene had descendants in more than one BUSCO orthogroup, we chose the orthogroup with highest overlap. We then computed the number of BUSCOs matched to a single ancestral gene, to two or more ancestral genes (dubious duplication), and absent from the ancestral genome reconstructed by AGORA (missing gene). Independent ancestral gene sets were downloaded from Ancestral Genomes (AG)^32^, based on PANTHER v.13.1^42^. Because AG and AGORA use different sets of extant species, we only considered ancestral genes with descendants in one of their common species for comparison (human for all ancestors except *Murinae* and *Laurasiatheria* where mouse and dog were used, respectively). AG ancestral genes were converted from UniProtKB IDs to Ensembl gene IDs using the correspondence tables provided by Ensembl BioMart, and compared with the gene sets in the ancestral genomes reconstructed by AGORA.

### Comparison of AGORA reconstruction with DESCHRAMBLER Eutherian ancestors

We compared AGORA’s v.92 eutherian reconstructions to DESCHRAMBLER’s (300 kb resolution: `APCF_hg19_merged.map` from http://bioinfo.konkuk.ac.kr/DESCHRAMBLER/). Since DESCHRAMBLER uses segments of the human genome as units of the reconstruction and was based on the hg19 genome assembly, we converted those regions to their proteincoding gene content, and selected the genes still found in Ensembl v.92 and descendants of ancestral Boreoeutherian genes. The Oxford grid plot were generated with the AGORA `src/misc.compareGenomes.py script in ‘matrix’` mode.

### Vertebrate evolutionary dynamics

Ancestral genomes reconstructed by AGORA from Ensembl version 102 were filtered to retain the most contiguous reconstructions, resulting in 73 ancestral genomes with G50 > 230 and L70 < 40. Conserved syntenic blocks between successive ancestral genomes in internal branches, and between ancestral genomes and their extant descendant in terminal branches were computed with PhylDiag^60^. Ends-of-blocks corresponding to likely evolutionary breakpoints were identified using *ad hoc* scripts. Orthologous genes between successive genomes were also compared in terms of their assignation to scaffolds or chromosomes larger than 200 genes using `AGORA’s src/misc.compareGenomes.py` script in `‘printOrthologousChrom’` mode. Groups of at least 20 genes relocating to more than one chromosome in a descendant genome, and inversely groups of at least 20 genes from two or more ancestral chromosomes relocating on the same descendant chromosome were considered inter-chromosomal rearrangements. Breakpoint and rearrangement rates per million years were computed using branch length estimates from TimeTree^61^. A full description of the parameters and selection thresholds are provided in Supplementary Information.

## Data availability

The source code of AGORA, user instructions and a test dataset are available for download from https://github.com/DyogenIBENS/Agora. Ancestral genomes have been precomputed for ~200 vertebrate (depending on the release), 41 plant, and 222 fungi genomes and are available on the Genomicus data base FTP server (ftp://ftp.bio.ens.psl.eu/pub/dyogen/genomicus/). These ancestral genomes can also be explored visually within the Genomicus^35^ synteny browser (http://www.genomicus.bio.ens.psl.eu/genomicus). Ancestral genomes used in this paper for analysis are archived on a Zenodo depository (https://sandbox.zenodo.org/record/962110).

## Acknowledgements

We thank Pierre Vincens for the coordination of computing resources and Amélie Peres for computer code engineering. This work was supported by grants from the French Government and implemented by ANR (ANR-07-GANI-008-01 GENOVERT, ANR-10-BINF-01-03 ANCESTROME, ANR-10-LABX-54 MEMOLIFE and ANR-10-IDEX-0001-02 PSL* Research University).

## Notes

https://www.genomicus.bio.ens.psl.eu/genomicus

https://github.com/DyogenIBENS/Agora

https://sandbox.zenodo.org/record/962110

### Competing Interest Statement

The authors have declared no competing interest.

## References

1. Zuckerkandl, E. & Pauling, L. Molecules as documents of evolutionary history. J. Theoret. Biol. 8, 357–366 (1965).

2. Felsenstein, J. Evolutionary trees from DNA sequences: a maximum likelihood approach. J Mol Evol 17, 368–376 (1981).

3. Yang, Z. Phylogenetic analysis using parsimony and likelihood methods. J Mol Evol 42, 294–307 (1996).

4. Nei, M. & Kumar, S. Molecular Evolution and Phylogenetics. (Oxford University Press, 2000).

5. Suzuki, Y. & Gojobori, T. A method for detecting positive selection at single amino acid sites. Molecular Biology and Evolution 16, 1315–1328 (1999).

6. Yang, Z. & Nielsen, R. Mutation-Selection Models of Codon Substitution and Their Use to Estimate Selective Strengths on Codon Usage. Molecular Biology and Evolution 25, 568–579 (2008).

7. Lupski, J. R. & Stankiewicz, P. Genomic disorders: molecular mechanisms for rearrangements and conveyed phenotypes. PLoS Genet 1, e49 (2005).

8. Rowley, J. D. Letter: A new consistent chromosomal abnormality in chronic myelogenous leukaemia identified by quinacrine fluorescence and Giemsa staining. Nature 243, 290–293 (1973).

9. Lupianez, D. G. et al. Disruptions of topological chromatin domains cause pathogenic rewiring of gene-enhancer interactions. Cell 161, 1012–25 (2015).

10. Tawn, E. J. & Earl, R. The frequencies of constitutional chromosome abnormalities in an apparently normal adult population. Mutat Res 283, 69–73 (1992).

11. Spielmann, M., Lupiáñez, D. G. & Mundlos, S. Structural variation in the 3D genome. Nat Rev Genet 19, 453–467 (2018).

12. Despang, A. et al. Functional dissection of the Sox9–Kcnj2 locus identifies nonessential and instructive roles of TAD architecture. Nat Genet 51, 1263–1271 (2019).

13. Schultz, J. & Dobzhansky, T. The Relation of a Dominant Eye Color in Drosophila Melanogaster to the Associated Chromosome Rearrangement. Genetics 19, 344–64 (1934).

14. Wilson, A. C., Sarich, V. M. & Maxson, L. R. The Importance of Gene Rearrangement in Evolution: Evidence from Studies on Rates of Chromosomal, Protein and Anatomical Evolution. Proceedings of the National Academy of Sciences 71, 3028–3030 (1974).

15. Sturtevant, A. H. Genetic factors affecting the strength of linkage in Drosophila. Proceedings of the National Academy of Sciences of the United States of America 3, 555–558 (1917).

16. Dobzhansky, T. & Sturtevant, A. H. Inversions in the Chromosomes of Drosophila Pseudoobscura. Genetics 23, 28–64 (1938).

17. Joron, M. et al. Chromosomal rearrangements maintain a polymorphic supergene controlling butterfly mimicry. Nature 477, 203–206 (2011).

18. Lowry, D. B. & Willis, J. H. A Widespread Chromosomal Inversion Polymorphism Contributes to a Major Life-History Transition, Local Adaptation, and Reproductive Isolation. PLOS Biology 8, e1000500 (2010).

19. Muñoz, A. & Sankoff, D. Detection of gene expression changes at chromosomal rearrangement breakpoints in evolution. BMC Bioinformatics 13 Suppl 3, S6 (2012).

20. M. Real, F. et al. The mole genome reveals regulatory rearrangements associated with adaptive intersexuality. Science 370, 208–214 (2020).

21. Loveland, J. L. et al. Functional differences in the hypothalamic-pituitary-gonadal axis are associated with alternative reproductive tactics based on an inversion polymorphism. Hormones and Behavior 127, 104877 (2021).

22. Lewin, H. A. et al. Earth BioGenome Project: Sequencing life for the future of life. PNAS 115, 4325–4333 (2018).

23. Armstrong, J. et al. Progressive Cactus is a multiple-genome aligner for the thousand-genome era. Nature 587, 246–251 (2020).

24. Yancopoulos, S., Attie, O. & Friedberg, R. Efficient sorting of genomic permutations by translocation, inversion and block interchange. Bioinformatics 21, 3340–3346 (2005).

25. Avdeyev, P., Jiang, S., Aganezov, S., Hu, F. & Alekseyev, M. A. Reconstruction of ancestral genomes in presence of gene gain and loss. 18.

26. Chauve, C., Gavranovic, H., Ouangraoua, A. & Tannier, E. Yeast Ancestral Genome Reconstructions: The Possibilities of Computational Methods II. Journal of Computational Biology 17, 1097–1112 (2010).

27. Tannier, E., Zheng, C. & Sankoff, D. Multichromosomal median and halving problems under different genomic distances. BMC Bioinformatics 10, 120 (2009).

28. Avdeyev, P., Jiang, S. & Alekseyev, M. A. Linearization of Median Genomes Under the Double-Cut-and-Join-Indel Model. Evol Bioinform Online 15, 1176934318820534 (2019).

29. Vakirlis, N. et al. Reconstruction of ancestral chromosome architecture and gene repertoire reveals principles of genome evolution in a model yeast genus. Genome Research 26, 918–32 (2016).

30. Chauve, C. & Tannier, E. A methodological framework for the reconstruction of contiguous regions of ancestral genomes and its application to Mammalian genomes. PLoS Comp Biol 4, e1000234 (2008).

31. Ma, J. et al. Reconstructing contiguous regions of an ancestral genome. Genome Research 16, 1557–1565 (2006).

32. Huang, X. et al. Ancestral Genomes: a resource for reconstructed ancestral genes and genomes across the tree of life. Nucleic Acids Res 47, D271–D279 (2019).

33. Kim, J. et al. Reconstruction and evolutionary history of eutherian chromosomes. Proceedings of the National Academy of Sciences 114, E5379–E5388 (2017).

34. Muffato, M., Louis, A., Poisnel, C. E. & Roest Crollius, H. Genomicus: a database and a browser to study gene synteny in modern and ancestral genomes. Bioinformatics 26, 1119–21 (2010).

35. Nguyen, N. T. T., Vincens, P., Dufayard, J. F., Roest Crollius, H. & Louis, A. Genomicus in 2022: comparative tools for thousands of genomes and reconstructed ancestors. Nucleic Acids Res gkab1091 (2021) doi:10.1093/nar/gkab1091.

36. Nguyen, N. T. T., Vincens, P., Roest Crollius, H. & Louis, A. Genomicus 2018: karyotype evolutionary trees and on-the-fly synteny computing. Nucleic acids research 46, D816–D822 (2018).

37. Boore, J. L. The use of genome-level characters for phylogenetic reconstruction. Trends Ecol Evol 21, 439–446 (2006).

38. Rokas, null & Holland, null. Rare genomic changes as a tool for phylogenetics. Trends Ecol Evol 15, 454–459 (2000).

39. Drillon, G., Champeimont, R., Oteri, F., Fischer, G. & Carbone, A. Phylogenetic Reconstruction Based on Synteny Block and Gene Adjacencies. Mol Biol Evol 37, 2747–2762 (2020).

40. Zhao, T. et al. Whole-genome microsynteny-based phylogeny of angiosperms. Nat Commun 12, 3498 (2021).

41. Manni, M., Berkeley, M. R., Seppey, M., Simão, F. A. & Zdobnov, E. M. BUSCO Update: Novel and Streamlined Workflows along with Broader and Deeper Phylogenetic Coverage for Scoring of Eukaryotic, Prokaryotic, and Viral Genomes. Molecular Biology and Evolution 38, 4647–4654 (2021).

42. Mi, H. et al. PANTHER version 16: a revised family classification, tree-based classification tool, enhancer regions and extensive API. Nucleic Acids Res 49, D394–D403 (2021).

43. Blanchette, M., Green, E. D., Miller, W. & Haussler, D. Reconstructing large regions of an ancestral mammalian genome in silico. Genome Research 14, 2412–23 (2004).

44. Ferguson-Smith, M. A. & Trifonov, V. Mammalian karyotype evolution. Nat Rev Genet 8, 950–62 (2007).

45. Stanyon, R., Stone, G., Garcia, M. & Froenicke, L. Reciprocal chromosome painting shows that squirrels, unlike murid rodents, have a highly conserved genome organization. Genomics 82, 245–249 (2003).

46. Murat, F. et al. Ancestral grass karyotype reconstruction unravels new mechanisms of genome shuffling as a source of plant evolution. Genome Research 20, 1545–57 (2010).

47. Romanov, M. N. et al. Reconstruction of gross avian genome structure, organization and evolution suggests that the chicken lineage most closely resembles the dinosaur avian ancestor. BMC Genomics 15, 1060 (2014).

48. O’Connor, R. E. et al. Reconstruction of the diapsid ancestral genome permits chromosome evolution tracing in avian and non-avian dinosaurs. Nature communications 9, 1883 (2018).

49. Jaillon, O. et al. Genome duplication in the teleost fish Tetraodon nigroviridis reveals the early vertebrate proto-karyotype. Nature 431, 946–57 (2004).

50. Carbone, L. et al. Gibbon genome and the fast karyotype evolution of small apes. Nature 513, 195–201 (2014).

51. Beklemisheva, V. R. et al. The Ancestral Carnivore Karyotype As Substantiated by Comparative Chromosome Painting of Three Pinnipeds, the Walrus, the Steller Sea Lion and the Baikal Seal (Pinnipedia, Carnivora). PLoS One 11, e0147647 (2016).

52. Pereira, A. L. et al. Extensive Chromosomal Reorganization in the Evolution of New World Muroid Rodents (Cricetidae, Sigmodontinae): Searching for Ancestral Phylogenetic Traits. PLoS One 11, e0146179 (2016).

53. Romanenko, S. A. et al. Multiple intrasyntenic rearrangements and rapid speciation in voles. Sci Rep 8, 14980 (2018).

54. Rhie, A. et al. Towards complete and error-free genome assemblies of all vertebrate species. Nature 592, 737–746 (2021).

55. Armstrong, J., Fiddes, I. T., Diekhans, M. & Paten, B. Whole-Genome Alignment and Comparative Annotation. Annual review of animal biosciences 7, 41–64 (2019).

56. Cunningham, F. et al. Ensembl 2022. Nucleic Acids Res gkab1049 (2021) doi:10.1093/nar/gkab1049.

57. Yates, A. D. et al. Ensembl Genomes 2022: an expanding genome resource for non-vertebrates. Nucleic Acids Res gkab1007 (2021) doi:10.1093/nar/gkab1007.

58. Peres, A. & Roest Crollius, H. Improving duplicated nodes position in vertebrate gene trees. BMC Bioinformatics 16, (2015).

59. Zdobnov, E. M. et al. OrthoDB in 2020: evolutionary and functional annotations of orthologs. Nucleic Acids Res 49, D389–D393 (2021).

60. Lucas, J. M., Muffato, M. & Roest Crollius, H. PhylDiag: identifying complex synteny blocks that include tandem duplications using phylogenetic gene trees. BMC Bioinformatics 15, 268 (2014).

61. Kumar, S., Stecher, G., Suleski, M. & Hedges, S. B. TimeTree: A Resource for Timelines, Timetrees, and Divergence Times. Molecular Biology and Evolution 34, 1812–1819 (2017).

