## Supplementary Information for "Reconstruction of hundreds of reference ancestral genomes across the eukaryotic kingdom"

[*] First authors

[†] Corresponding authors

#### Supplementary Material

#### Glossary

Here is a summary of the AGORA-specific terms we use in this document:

- Ancestral gene — Inferred existence of a gene in a given ancestor.
- Constrained ancestral gene — Ancestral gene that has undergone fewer duplications and losses than a given threshold.
- Orthology group — Set of extant genes that derive from a single speciation event. AGORA represents ancestral genes as orthology groups.
- Gene pair — Any two genes.
- Gene adjacency — Two contiguous genes on a chromosome, taking their transcriptional orientation into account.
- Conserved gene adjacency — Gene adjacency that is seen in two or more genomes, using orthologues to do the comparison.
- Contiguous ancestral region — (abbreviated as CAR). Ordered list of oriented ancestral genes, representing the region of an ancestral genome. A CAR may correspond to an entire chromosome in an ancestral genome, or a portion of it.
- Singleton — Ancestral gene that could not be placed in any CAR.
- Ancestral genome — A collection of CARs and singletons for a given ancestor, that encompasses all its ancestral genes.
- Ancestral block — Output of a reconstruction step. A block is an ordered list of oriented elements, which can be either ancestral genes or ancestral blocks.
- Integration — A workflow step that builds ancestral blocks from conserved adjacencies.
- Single-integration pass — Reconstruction workflow that considers all ancestral genes (or blocks) at once.
- Multi-integration pass — Reconstruction workflow that processes constrained and non-constrained ancestral genes (or blocks) differently across multiple integration steps.
- One-pass reconstruction — Reconstruction workflow that runs a single pass.
- Two-pass reconstruction — Reconstruction workflow that runs two passes. The blocks reconstructed during the second pass are made of the blocks reconstructed during the first pass.

#### AGORA method

##### *Overview*

The AGORA method (outlined in Fig. S1) is a generic and flexible framework for reconstructing ancestral genomes by comparing extant genomes. The rationale that underlies AGORA is that similarities between any two genomes often reflect ancestral features that existed in all the ancestors that lie on the evolutionary path that leads from one genome to the other in the species tree. To distinguish the similarities that occur by chance from those that truly reflect an ancestral state, AGORA integrates data over tens to thousands of comparisons for a given ancestral genome to accumulate confidence in the selected similarities.

AGORA requires two sets of information to reconstruct ancestral genomes: the position of genes in their respective extant genomes and the phylogenetic relationships among these genes. This data is processed by AGORA in several steps: (i) ancestral gene content extraction if not provided by the user, (ii) pairwise comparisons and (iii) integration. AGORA does not annotate genomes and does not compute phylogenetic gene trees, both of which must be provided by the user. Input gene trees have to be reconciled with the species phylogeny. While a range of resources exists to obtain or to compute such data, we used the Ensembl[1] database as a central source of homogeneous and exhaustive information on both gene annotations and gene phylogenetic trees.

##### *Extraction of ancestral genes*

For a given gene, a reconciled phylogenetic tree records the complete history of its evolution, including speciation, duplication and loss. Ancestral genes can thus be inferred from the gene trees. It is possible to establish the gene catalogue of all ancestors by traversing the complete set of phylogenies and adding genes to the relevant ancestors.

AGORA represents an ancestral gene as an orthology group: the set of extant genes that derive from it, as per the gene tree (Fig. S2). Orthology groups can be provided by the user, or will be inferred from the gene trees by AGORA. Ancestral genes of each ancestor are identified by traversing each gene tree from its root node thanks to the reconciliation tags. When a duplication node is encountered, AGORA creates an additional ancestral gene at this duplication node’s ancestor, and splits the extant gene content across both ancestral genes. When a speciation node is encountered or when AGORA hits a leaf, AGORA marks the ancestral gene as present on all ancestors since the last one it considered, as it does between two consecutive duplication events that happened in different ancestors. Finally, when a loss event is encountered, AGORA stops marking ancestral genes in this lineage.

Within this framework, when an extant gene does not relate to any genes of a given ancestor, it is considered a lineage-specific creation (relative to that ancestor). This can happen if the root of its gene tree is younger than the ancestor considered. Two extant genes are orthologous if they descend from the same ancestral gene of their last common ancestor. Two extant genes are paralogous if they descend from different ancestral genes of their last common ancestor but the same ancestral gene of an older ancestor. Two extant genes are not homologous if no ancestral gene relates to both.

##### *Selection of constrained ancestral genes*

A reasonable assumption is that an ancestor's gene set must have been similar in size to the extant species below it (a notable exception is clades that underwent a whole genome duplication). As reported in Figure 3A, AGORA initially tends to overestimate the number of genes present in each ancestral genome. This behavior is due to the existence of poorly supported duplication nodes in the gene trees, leading the algorithm to infer the existence of two paralogous copies of a gene in the ancestral genome. Over 40% of duplication nodes in the gene trees of Ensembl have a “duplication confidence score”[2] lower than 0.30 (86,315 duplication nodes in total as of database v.85). Erroneous duplication nodes result in the inference of a 'phantom' gene copy in the ancestor. However, these 'phantom' genes have only one or few descendants in modern genomes (hence their dubious duplication status) and usually remain singletons in the AGORA reconstructions due to lack of placement support.

AGORA overcomes this by identifying a subset of ancestral genes that are coined as “constrained”. These ancestral genes are represented by a number of extant genes close to the number of species that descent from the ancestor, thus having undergone few duplications and losses. For vertebrates, that range is defined as a gene-to-species ratio within 90%-110%. Such ancestral genes are more uniformly annotated and provide a clearer picture when comparing the species.

AGORA is able to employ a multi-step strategy to leverage these constrained genes, by first building backbones of ancestral genomes using constrained gene families only, and then reconstructing local gene order using the remaining gene families.

##### *Pairwise comparisons – extraction of conserved gene adjacencies*

The core principle of this step is that when two genes are consecutive in one genome and their two respective orthologues are also consecutive and in the same transcriptional orientation in another genome, then the ancestral copies of these genes probably existed in the same configuration in all the ancestral nodes between the two species, from their last common ancestor. This definition of conservation, combining strict adjacency and transcriptional orientation of two functional sequences, is very stringent and unlikely to occur by chance.

Depending on its position in the tree, an ancestral genome may be assigned a conserved adjacency through different comparisons. Indeed, an ancestral node is always found at the cross road of three branches: two descendants and one outgroup (except for the root of the species tree). Any comparison between two species that belong to two of the three branches is potentially informative to identify a conserved adjacency in that ancestor.

The naive implementation would result in a $O(n^2 \times \log(n))$ time complexity ($n^2$ comparisons, and $\log(n)$ ancestors to propagate the conserved adjacencies to). AGORA implements an efficient algorithm to perform all the comparisons in a $O(n \times \log(n))$ time complexity at the expense of memory, by precomputing data into hash tables.

First, all extant genomes are iteratively filtered down to the gene set of each of their ancestors, and all the gene adjacencies are extracted (Fig. S3). For each ancestor *A*, AGORA considers all the pairs of species *below* it that are under different children, takes their set of gene adjacencies filtered down to that ancestor *A*, and computes the pairwise intersection. AGORA takes the union of all those intersections, while counting how many comparisons have contributed to each conserved adjacency. Finally, it marks each of these adjacencies as conserved in the ancestors that lie between the ancestor *A* and the extant species that contributed to it, unless they are disrupted by a recent insertion.

For each ancestor, the result is a list of oriented gene adjacencies with the number of comparisons that support it. This step is performed twice: on the set of constrained gene families and on the complete set.

##### *Integrations – ancestral genome reconstructions*

Hereafter, the AGORA steps are called “integrations” as they combine in various ways the conserved adjacencies that have been identified in order to generate ancestral genomes. The “de novo” integration reconstructs ancestral genomes solely using the conserved adjacencies, whereas the other integration steps also use the output of a previous integration step. The fundamental rule that prevails is that each further integration step preserves the relative order of the genes that are in its input reconstructions. Reconstructions incrementally grow and provide a more and more complete view of the ancestral genomes.

AGORA is typically used in two settings: a single-integration mode, where only the “de novo” integration is run, and a multi-integration mode, where AGORA runs the “de novo” algorithm on the set of constrained ancestral genes, and then other integration steps in order to add the non-constrained genes.

##### *“De novo” integration*

Here, for each ancestor, the conserved adjacencies identified by the algorithm above are represented in a weighted adjacency graph where nodes are oriented ancestral genes, edges represent observed conserved adjacencies, and the weights are the number of comparisons that support each adjacency (Fig. S4). In a perfect scenario, the graph would be acyclic, with node degrees no greater than 2, thus immediately providing the structure of the ancestral genome as chromosomes. In reality, the high number of pairwise comparisons identifies a large amount of conserved gene adjacencies, which are not always consistent between each other because of evolutionary rearrangements, assembly, annotation or gene tree reconstruction errors, or evolutionary convergence. This results in the graph usually containing cycles and bifurcations.

AGORA employs a greedy strategy to partition the graph into a set of acyclic, non-overlapping paths, selecting the highest weighted edges first, and adding edges of lower weight as long as they do not create forks or cycles with the previously selected edges.

The result is a set of contiguous oriented genes (similar to contigs in a sequence assembly process) that represent ancestral chromosomes (or portions of chromosomes) and a set of singletons. The blocks cannot be extended on either side because the genes at the ends either are not involved in any conserved adjacency (i.e. have different neighbours in all the genomes tested in the pairwise comparisons), or their adjacencies contradict other blocks. The same reason holds for singletons (blocks of length 1).

##### *“Fill-in” integration*

In this step, AGORA fills the blocks created by the “de novo” integration with non-constrained genes, using the conserved adjacencies identified when comparing all the genomes on the entire sets of ancestral genes.

This is done by representing the conserved adjacencies in weighted adjacency graphs that are anchored into the blocks (Fig. S5). The graphs themselves (nodes, edges, weights) are constructed the same way as in the “de novo” step. AGORA searches paths of non-constrained genes that link consecutive constrained genes and do not create cycles nor bifurcations. AGORA seeks to select the longest possible paths in order to maximise the number of genes included in the reconstructions, but those paths may contradict each other. For instance, an ancestral gene may be part of the longest paths of two different intervals. In such cases, AGORA chooses the paths that has the highest sum of weights along the adjacencies that it includes, and discards the other. AGORA then tests the next longest path for the second interval.

This iterative process results in each interval of constrained genes being filled with 0 or more non-constrained genes (ordered and oriented). All such extensions are compatible with one another (no cycles, no bifurcations). The output contains an additional set of singletons: the non-constrained genes that could not be added to any interval.

##### *“Fusion” integration*

In this step, AGORA applies the “de novo” algorithm on the singletons, which contain both constrained and non-constrained ancestral genes (Fig. S6). Although the constrained singleton genes are not part of any conserved adjacencies between themselves (otherwise they would have formed a block in the first “de novo” step), they may be involved in conserved adjacencies with non-constrained genes. Non-constrained genes can also be part of conserved adjacencies between themselves.

The output is a set of additional blocks that replace the singleton genes they are made of.

##### *“Insertion” integration*

At this stage, all the gene adjacencies within the blocks reconstructed are conserved. Moreover, since the “fill in” algorithm does a *longest* path search, the blocks cannot be extended without breaking that property of the blocks.

The “insertion” step (Fig. S7) acknowledges that errors in genome assembly, annotation and gene tree reconstruction can happen, and result in accidental loss of gene order conservation. In this step, AGORA seeks to insert the blocks created in the “fusion” integration step into the blocks created in the “fill in” step, while requiring *only one* of their ends to be supported by a conserved adjacency. The latter must have a higher weight than the one that supports the target interval. The ratio of these two weights (which is higher than 1) is what AGORA uses to choose which insertions to perform. Each interval *A*-*B* can welcome two insertions: on the right side of *A* and on the left side of *B*.

As in the “de novo” step, AGORA employs a greedy strategy to select the insertions, considering the ones with the highest weight ratios first, and then the ones with lower weight ratios as long as they do not target the same insertion point. AGORA also tries to extend the blocks on each of their ends, using the weights to rank the possible extensions. A “fusion” block cannot be inserted in more than one point.

In the resulting blocks, not every adjacency is conserved amongst extant genomes, but non-conserved adjacencies are always surrounded by conserved ones, and within large-scale gene order conservation blocks.

##### *Blocks of blocks – two-pass reconstruction*

In the same way that extant genomes have been considered as sequences of oriented genes, and the ancestral order of these genes has been reconstructed into blocks, extant genomes can be described as sequences of blocks and the ancestral order of these blocks can be reconstructed. This is similar to the scaffolding in the genome sequence assembly process. The method described below is an adaptation of the gene-based method presented above, working on blocks instead of genes, and forms the second pass of a two-pass reconstruction. Although the same methods could be used for a third pass (blocks of blocks of blocks) and more, we only use two passes for all our reconstructions.

##### *Pairwise comparisons – extraction of conserved block adjacencies*

The adjacency measured between the blocks has to be more relaxed than the one between the genes. The difficulty lies in the fact that a reconstructed block is not necessarily continuous in each extant genome, but perhaps interrupted by rearrangements. Indeed, since blocks are the result of integration steps of all conserved adjacencies between all genomes, a block can include two regions of different chromosomes of an extant species.

To identify block adjacencies, we need to identify the position of the extremities of the ancestral blocks on extant genomes, and then extract the cases where ends of different blocks are contiguous. AGORA starts by “aligning” the ancestral blocks with the extant genomes: identifying sequences of consecutive genes (at least two) that are in the same order and same transcriptional orientation. An adjacency between two blocks is declared when the first aligned segment of one block immediately follows the last aligned segment of the other block in an extant genome (respecting the transcriptional orientations). AGORA actually considers all possible relative orientations when searching block adjacencies, i.e. C1 followed by C2 (both in their default orientation), C1 followed by the reverse of C2 in its opposite orientation, etc.

For a given ancestor, AGORA builds the set of block adjacencies of each extant genome. Then it compares every pair of descendants that are under different children, and every descendant to every outgroup to intersect their respective sets of block adjacencies and build a weighted adjacency graph, where the weight is the number of comparisons that support the adjacency.

##### *Integration overview*

The “de novo”, “fill-in”, “fusion”, and “insertion” algorithms described above can be used to build blocks of blocks. We typically use the single-integration mode (i.e. only the “de novo” algorithm) for Vertebrates, and the multi-integration mode for Plants.

##### *Multi-integration reconstruction*

To run a multi-integration reconstruction for Plants, we need to define a filter that marks some of the first pass blocks as “constrained”. We have experimented with several such filters but have not been able to reliably identify one that outperforms the others. Our Plants workflow hence runs four versions of the second pass using different filters, and then chooses, for each ancestor, the version that yields the highest G50 (see Methods). The four filters are:

- all the blocks of 20 genes or more
- all the blocks of 50 genes or more
- the longest blocks that encompass 50% of the ancestral genome
- the longest blocks that encompass 70% of the ancestral genome

#### Software implementation and packaging

AGORA is available as a set of Python scripts on GitHub at https://github.com/DyogenIBENS/Agora, licensed under the GNU General Public License version 3 (GPL v3) and the CeCILL licence version 2 of the CNRS.

AGORA has a small number of dependencies, and is compatible with the reference Python implementation (CPython) and PyPy versions 3 or above. An example containerisation as a Docker image is also provided.

AGORA comes with its own workflow manager to (i) extract the ancestral gene content, (ii) do the pairwise comparisons, and (iii) run the integration steps (single-integration and multi-integration, one or two passes). The most common scenarios are directly available through these scripts:

- `agora-basic.py` runs two single-integration passes (black path on Fig. S1). This is the first script to try on a dataset.
- `agora-generic.py` runs two multi-integration passes, and should be used when the `agora-basic.py` reconstructions are too incomplete or fragmented. Each pass is run with several filters that select the “constrained” genes or blocks, and AGORA automatically selects the version with the highest G50. This workflow takes longer to execute than `agora-basic.py` but will output a more complete reconstruction.

- `agora-vertebrates.py` is the workflow used for the Vertebrates reconstructions (red path on Fig. S1).
- `agora-plants.py` is the workflow used for the Plants reconstructions (green path on Fig. S1).

AGORA takes standard file formats as inputs: reconciled gene trees in NHX, species tree in Newick, and gene content of the extant species as BED files. It outputs all its data (intermediate and final reconstructions) in tabular formats and the final ancestral genomes are also available as BED files.

#### Benchmarks against simulations

##### *Simulations from Kim et al., 2017 (DESCHRAMBLER)*

We applied AGORA (`agora-basic.py`) to the 50 simulated datasets from the DESCHRAMBLER publication[3]. In these datasets, the genomes are described as lists of oriented markers (1 list for each chromosome), in files named `SG_ALL_GENOMES` for the extant genomes, and `SG_ANCESTOR.{boreo,euarch,rodent}` for the ancestral genomes. First, the 9 simulated extant genomes were converted to the BED-like format required by AGORA. The ancestral "genes" were inferred from the markers' presence/absence patterns in the extant genome using parsimony, e.g. a marker only seen in human and mouse was listed in *Euarchontoglires*' ancestral genes set, but not *Boreoeutheria*'s. The reconstructed genomes, and the true, simulated, ones were converted to sets of adjacencies of oriented genes in order to compute precision, sensitivity, and the Jaccard index (called "agreement" in [2]). Like DESCHRAMBLER, AGORA achieves on average >99% precision and sensitivity on all three ancestors *Boreoeutheria*, *Euarchontoglires*, *Rodentia* (the standard deviation is indicated in parentheses).

| Ancestor | Precision | Sensitivity | Jaccard index |
|---|---|---|---|
| ***Boreoeutheria*** | 99.62% (0.06) | 99.30% (0.14) | 98.92% (0.15) |
| ***Euarchontoglires*** | 99.64% (0.06) | 99.83% (0.07) | 99.47% (0.10) |
| ***Rodentia*** | 99.56% (0.08) | 99.04% (0.13) | 98.61% (0.18) |

##### *Simulations using MagSimus*

A striking limitation of the simulations from the DESCHRAMBLER publication is that they do not model duplications despite these being ubiquitous in gene evolution. For instance, in the version 92 of Genomicus (based on Ensembl 92), out of the 22,773 inferred genes for *Boreoeutheria*, 19,463 are in multiple copies in at least one descendant species.

We generated our own, more realistic, simulations that feature a more complete set of rearrangement and events, including duplications. The simulator, named MagSimus, is available at https://github.com/DyogenIBENS/MagSimus, licensed under the GNU General Public License version 3 (GPL v3) and the CeCILL licence version 2 of the CNRS (like AGORA).

MagSimus[4] models genomes as lists of ordered markers that represent genes. Starting from the root of the species tree, an initial, random, genome undergoes random events that change its gene content (duplications, deletions, births) and gene order (inversions, translocations, fusions, fissions) successively on all branches of the species tree, towards the leaves. The rates of each type of event have been estimated from the actual genomes. Global rates are defined in the file named data/parametersG_ABC.v83, and per-branch rates in data/specRates_MS1.v84. The latter lists the rates of the seven events listed above, as well as the proportion of gene duplications that happen in tandem (as opposite to the new copy being inserted randomly in the genome). Importantly, translocations, fusions and fissions select the chromosomes they affect independently of their length whereas the gene events and inversions select genes and intervals at random, thus selecting chromosomes proportionally to their length. The inversion lengths follow a Gamma distribution of shape $k$=1 and scale $\theta$=21.3630. The consequence of those parameters is that the simulated *Boreoeutheria* genomes all had 24 chromosomes and 23,445 genes.

The output is a set of simulated extant genomes (human, mouse, dog, opossum, chicken, as in the previous simulation set, Figure S16), their ancestral genomes, and files similar to AGORA's "ancestral genes" (i.e. the evolution of the gene families). Those files all had to be reformatted to fit AGORA's and DESCHRAMBLER's input formats. Specifically, for AGORA, we prefixed all gene names with their species name to make them unique. For DESCHRAMBLER, we converted the orthology groups to "conserved regions", after discarding (i) duplications, (ii) the ones that are not single copy in human, (ii) the ones that are not single copy in any other species.

The first difference between both methods is that AGORA effectively operates on more genes than DESCHRAMBLER: on average 22,496.6 vs 11,045.3 (out of 23,445, over 50 iterations). While AGORA accepts all ancestral genes as inputs, 948.4 genes on average end up as singletons as they have less than two extant genes, or these are located within a single sub-tree attached to *Boreoeutheria*. For DESCHRAMBLER, the difference comes from the constraints of the "conserved regions" file explained above. This highlights the lack of resolution that results from only considering single-copy genes.

Then, despite being able to assemble a higher proportion of its usable genes into CARs, DESCHRAMBLER's reconstructions still only contain 9,900.4 genes on average versus 19,120.0 for AGORA, meaning the resolution of DESCHRAMBLER is about half that of AGORA.

| Method | Usable genes | Genes in CARs | Coverage | |
|---|---|---|---|---|
| | | | Usable genes | Whole genome |
| **AGORA** | 22,496.6 genes (25.7) | 19,120.0 genes (41.1) | 85.0% (0.20) | 81.6% (0.18) |
| **DESCHRAMBLER** | 11,045.3 genes (42.4) | 9,900.4 genes (53.4) | 89.6% (0.29) | 42.2% (0.23) |

Average coverage statistics after 50 MagSimus simulations, standard deviation in parentheses

Reconstructed adjacencies were compared against the original simulated ones considered as "truth", after singletons were excluded. Due to is ability to consider every gene from every species and not using a reference species, AGORA achieves 95.4% agreement, significantly higher than DESCHRAMBLER's (68.6%), while running 190 times faster.

| Method | Precision | Sensitivity | Agreement | Runtime (per simulation) |
|---|---|---|---|---|
| **AGORA** | 98.7% (0.11) | 96.6% (0.16) | 95.4% (0.24) | 20.0 sec. (0.5) |
| **DESCHRAMBLER** | 86.4% (0.61) | 76.9% (0.61) | 68.6% (0.86) | 3,799.0 sec. (238.5) |

Average reconstruction quality for 50 MagSimus simulations, standard deviation in parentheses

#### Vertebrate genome evolutionary dynamics.

**Data**

All analyses are based on genomes from Ensembl[1] version 102. This version references 310 genomes, of which 269 are used for the Ensembl Compara database. Ensembl Compara includes gene trees built using the TreeBest pipeline and made publicly available.
We started from this set of 269 genomes and removed the following subsets:

- 21 genomes produced by the Vertebrate Genome Project (VGP) for embargo issues,
- 152 genomes of low contiguity based on Extended Figure 10a,
- descendants of "*Eupercaria incertae sedis*" for formatting issues,

- non-vertebrate genomes, and
- descendants of 13 ancestral genomes of low contiguity based on Extended Figure 10b.

This filtering resulted in 74 extant species (15 birds and reptiles, 41 mammals, 18 teleostean fish) represented in a species tree with 146 branches connected by 73 ancestral nodes for which AGORA had reconstructed ancestral genomes (Figure 5a).

**Computing rearrangement breakpoints**

Rearrangement breakpoints are located at the edges of syntenic blocks. To compute syntenic blocks, we used PhylDiag[4] between all pairs of successive genomes found in the species tree (either ancestor-ancestor in internal branches, or ancestor-extant in terminal branches) with the following parameters:

```
phylDiag.py --no-imr -m 50 -t 5 -g 45
```

Careful examination of early results showed that false positive breakpoints were caused by ends-of-blocks that:

- represent extremities of scaffolds or chromosomes,
- are located in ancestral gene adjacencies that are not or poorly supported in the AGORA adjacency graph,
- are located in ancestral or extant scaffolds smaller than 10 genes, or
- are located within 3 genes of scaffold or chromosome ends.

Custom Perl and Python scripts were written to identify and exclude synteny block ends fulfilling these criteria. Next, because the ancestral state (pre-breakpoint) and the descendant state (post-breakpoints) are known, the resolution of synteny block ends into breakpoints is immediate, in the form of two ends-of-syntenic blocks that are adjacent in the ancestral genome. A custom Python script was used to count all such instances as breakpoints (Supplementary Table S2).

**Computing interchromosomal rearrangements**

To compute interchromosomal rearrangements, we compared the chromosome assignation of genes between two successive nodes in the tree on Figure 5 using AGORA's src/misc.compareGenomes.py utility, restricting the comparison of chromosomes containing at least 200 genes, with the following parameters:

```
src/misc.compareGenomes.py genome1 genome2 genome2 \
                           -mode=printOrthologousChrom \
                           -minChrSize=200
```

A custom Python script was then used to identify cases where genes (at least 20) from a chromosome in genome1 were distributed on more than one chromosome in genome2. Each group of at least 20 genes was counted (total = N) and the number of rearrangements was considered to be: rearrangements = N – 1. Similarly, the same script identified cases were groups of at least 20 genes residing on two different chromosomes in genome1 are located on the same chromosome in genome2. Each such case was counted as an additional rearrangement (Supplementary Table S2).

**Computing rates (Figure 5b)**

All branch lengths in million years were computed based on ancestral node ages provided by TimeTree[5] (Supplementary Table S3).

#### Supplementary Figures and Figure Legends

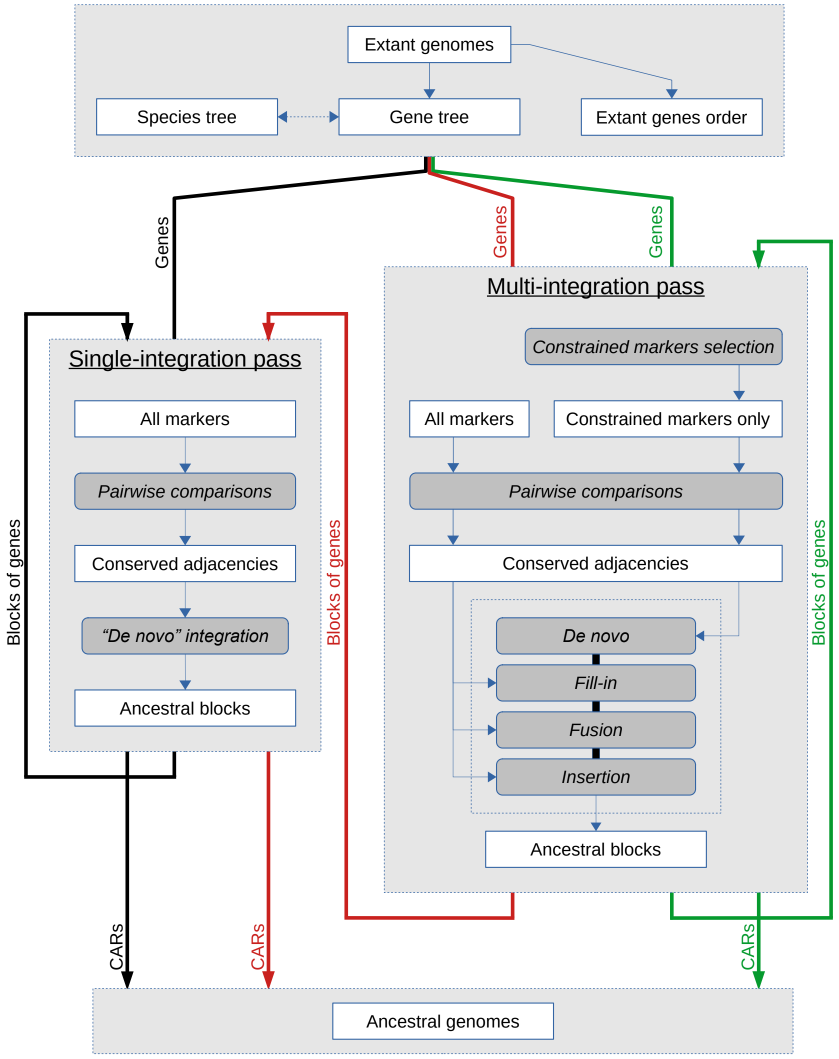

**Fig. S1**

Overview of the AGORA workflow. Input data are at the top, AGORA in the middle, and output data at the bottom. White rectangles represent data, grey rounded rectangles indicate processes (steps), which are arranged in two modules: single-integration pass and multi-integration pass. A reconstruction is a series of one or two passes (typically two). The thick black arrow indicates a basic workflow that consists of two single-integration passes, going from genes to blocks of genes (first pass) and to blocks of blocks of genes (second pass) that form the CARs of the ancestral genome. The red path shows a two-pass reconstruction that does a single-integration following a multi-integration (this is typically used for vertebrate genomes). The green path shows two consecutive multi-integration reconstructions, which is typically used for plant genomes.

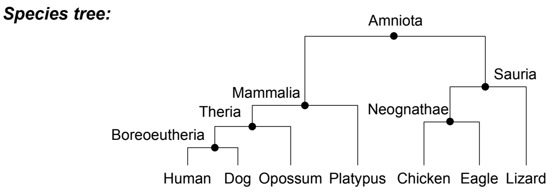

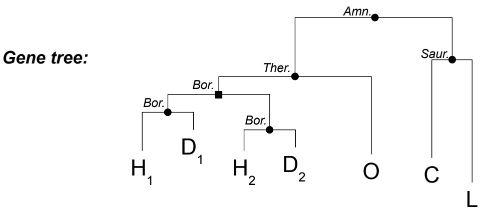

| Ancestor | Ancestral genes | Orthologues | Paralogues |
|---|---|---|---|
| **Amniota** | {C, L, O, $H_1$, $H_2$, $D_1$, $D_2$} | {C, L} – {O, $H_1$, $H_2$, $D_1$, $D_2$} | |
| **Sauria** | {C, L} | {C} – {L} | |
| **Neognathae** | {C} | | |
| **Mammalia** | {O, $H_1$, $H_2$, $D_1$, $D_2$} | | |
| **Theria** | {O, $H_1$, $H_2$, $D_1$, $D_2$} | {O} – {$H_1$, $H_2$, $D_1$, $D_2$} | |
| **Boreoeutheria** | {$H_1$, $D_1$} | {$H_1$} – {$D_1$} | {$H_1$, $D_1$} – {$H_2$, $D_2$} |
| | {$H_2$, $D_2$} | {$H_2$} – {$D_2$} | |

**Fig. S2**

Inference of ancestral genes, orthologues, and paralogues. The species tree is represented in the top panel as a cladogram, as only the species hierarchy and the clade names are needed. The gene tree (in the middle panel) is reconciled with the species tree: speciation nodes are drawn as black circles, duplication nodes as black squares, both being associated with a clade name. The table (in the bottom panel) lists the ancestral genes, the orthologues, and the paralogues. Ancestral genes are given as the set of extant genes that derive from it. Orthologues (resp. paralogues) are written as the cartesian product of two sets: each gene from the first set is orthologous (resp. paralogous) to every gene from the second set. The oldest ancestral gene that can be inferred sits at the *Amniota* ancestor and encompasses all known copies of this gene, including two copies in human and dog, but none in platypus and eagle. Every other ancestor (except *Boreoeutheria*) have a single ancestral gene too, linking to the extant copies that evolved from it. Ancestors that are not directly represented in the gene tree (*Neognathae* and *Mammalia*), e.g. because the extant genes are confined to only one of their child branches, can still have ancestral genes attached to them. Ancestral genes are also created when there is a single extant copy remaining (*Neognathae*). *Boreoeutheria* is the only ancestor with two ancestral genes (because of a gene duplication) which separate out the two human and dog copies of the gene.

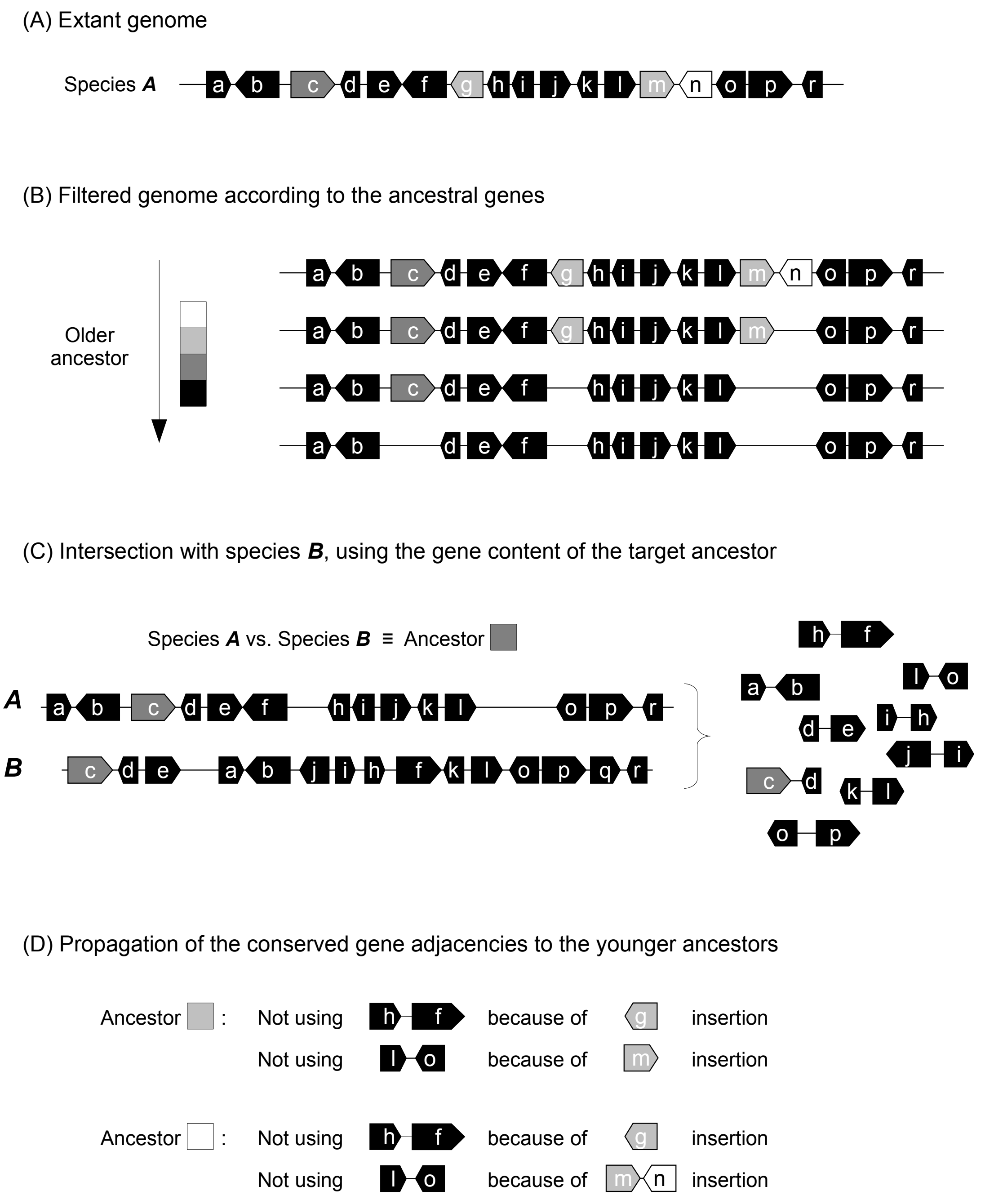

**Fig. S3**

Pairwise comparisons: extraction of conserved adjacencies. When comparing two species, AGORA maps their genomes to the ancestral gene content of their last common ancestor, extracts the ancestral gene adjacencies and computes the intersection of both sets. According to the principle of parsimony, conserved gene adjacencies are considered as being present in all the ancestors that lie on the evolutionary path, but AGORA needs to discard the ones that are interrupted by the apparition of (more recent) ancestral genes.

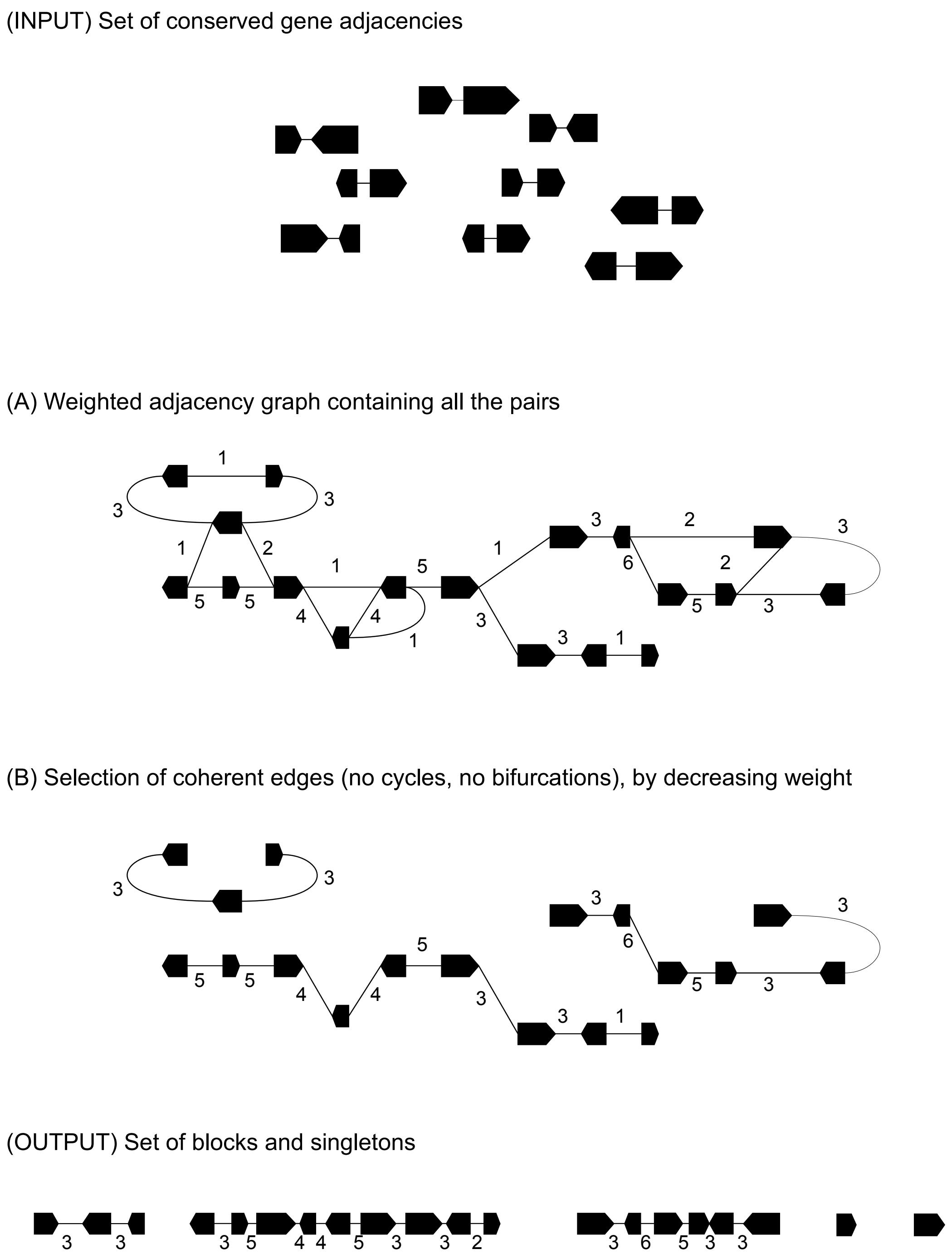

**Fig. S4**
"De novo" integration. For a given ancestor, all the conserved gene adjacencies are combined into a weighted directed graph, from which edges are selected by decreasing weight in order to make a subgraph that does not contain any cycle or bifurcations. Each connected component defines the relative order of some ancestral genes in the ancestral genome.

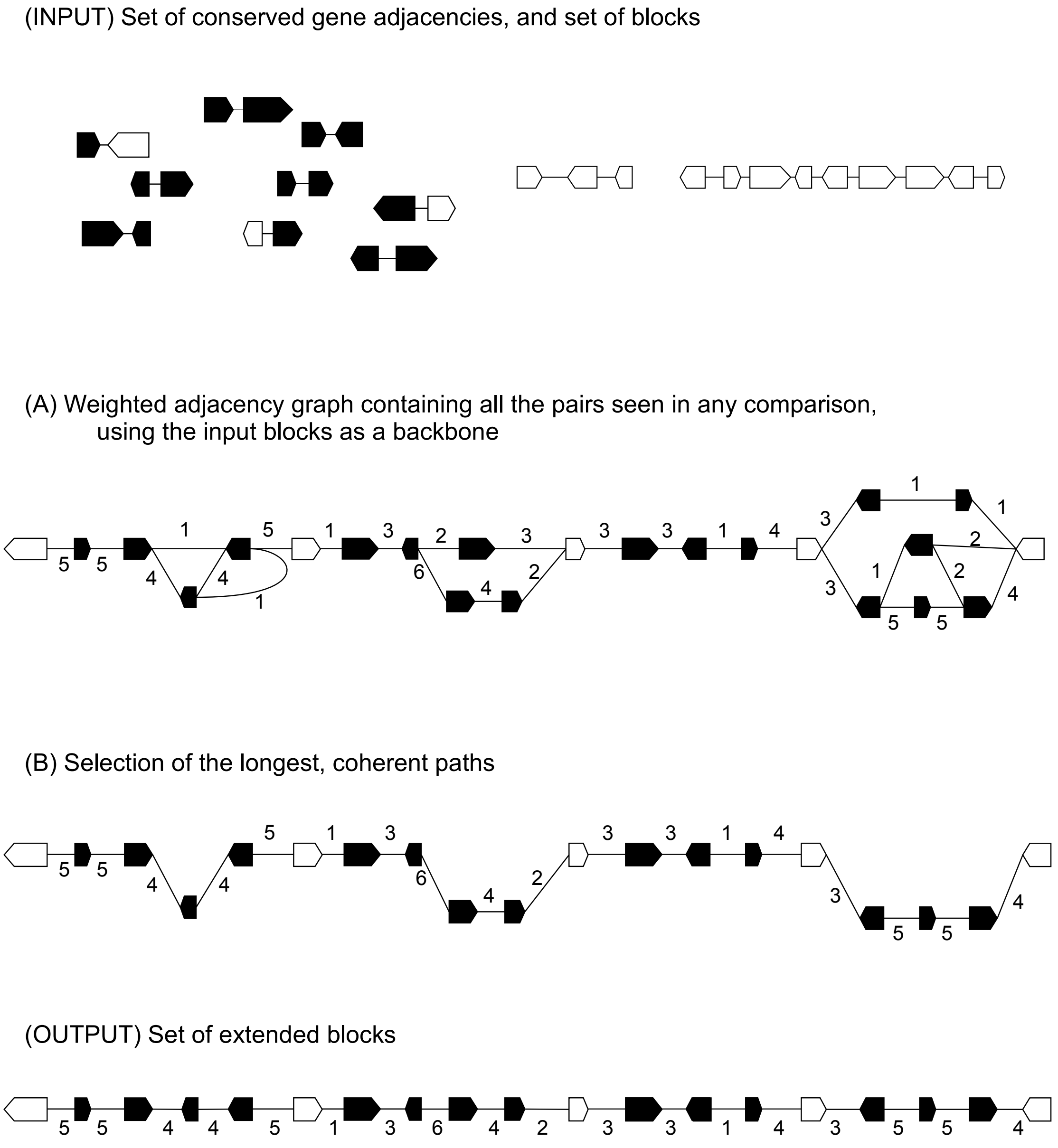

**Fig. S5**

<u>"Fill-in" integration</u>. For a given ancestor, all the conserved gene adjacencies are combined into a weighted directed graph anchored onto the blocks created by the "de novo" integration. AGORA selects the longest path within each adjacency of constrained genes in order to maximise the number of genes included in the reconstructions, as long as they do not conflict with other longest paths. The weights are used to resolve such conflicts.

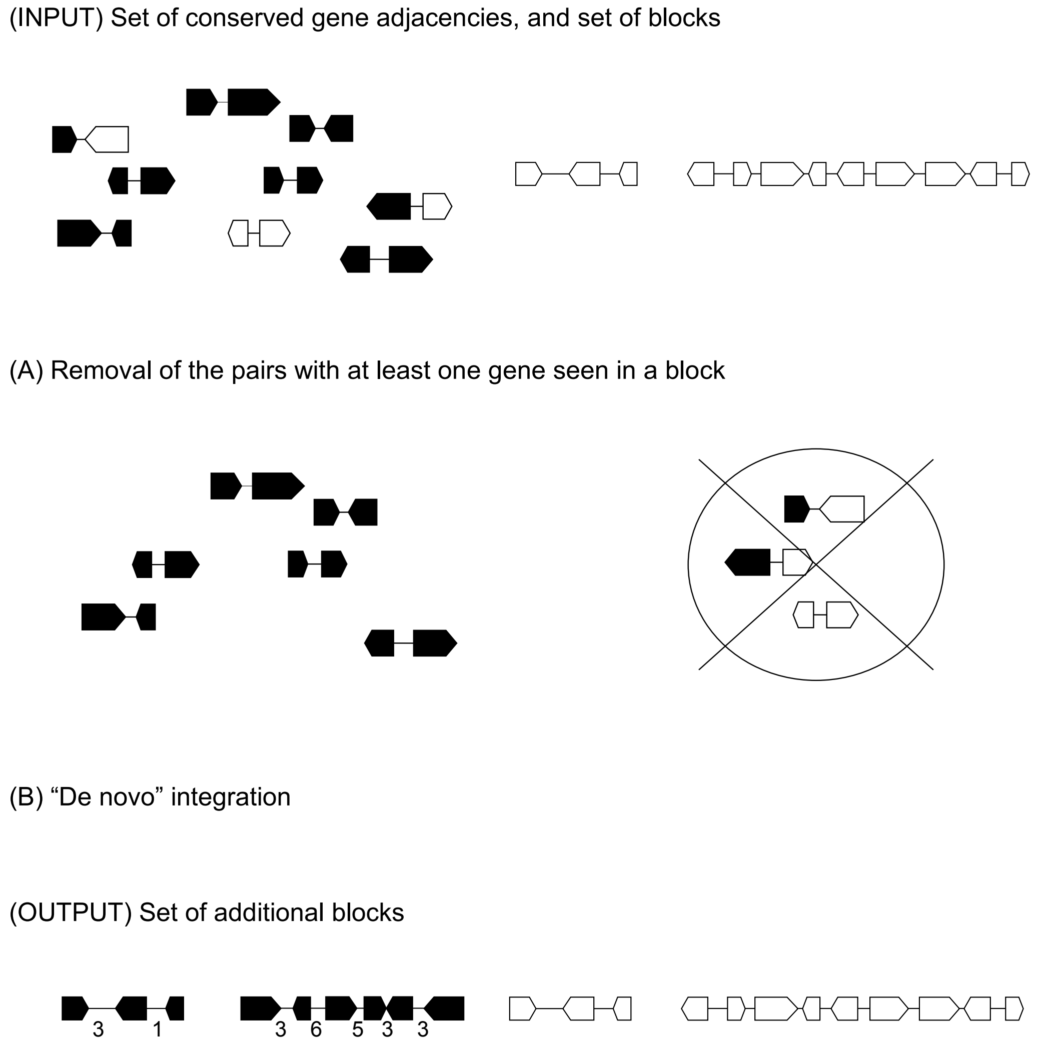

**Fig. S6**
<u>"Fusion" integration</u>. In this step, the singletons (constrained and non-constrained) have a chance to form new blocks, independently of the existing blocks, through the same process as a "de novo" integration.

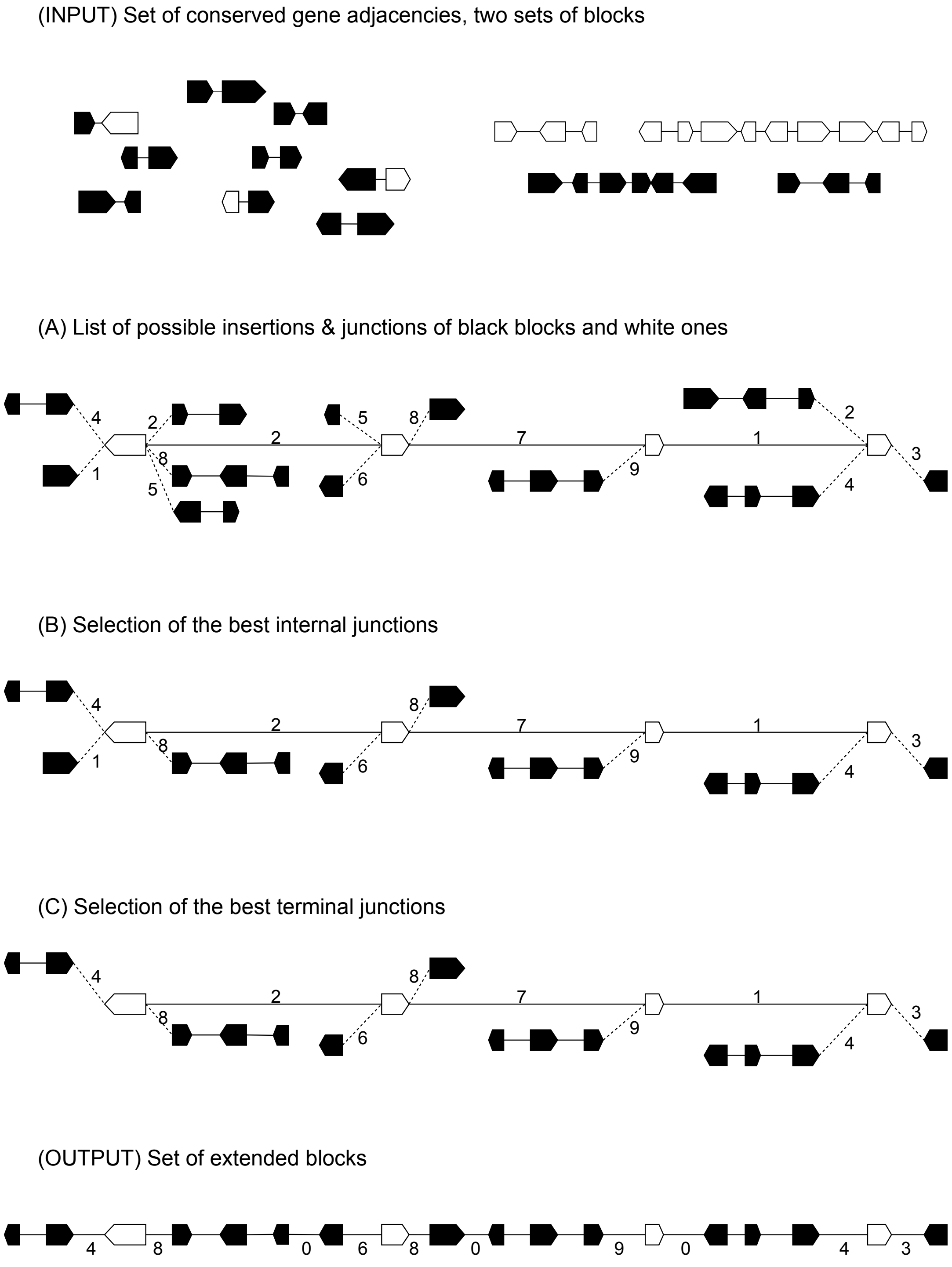

**Fig. S7**

<u>"Insertion" integration</u>. In this step, the blocks created during the "Fusion" integration will be inserted into the blocks from the "Fill-in" integration using observed conserved adjacency and considering the weights to solve conflicts. The insertions can in reality only be supported by one side, meaning that the resulting blocks feature some adjacencies that are not directly observed in any extant genome.

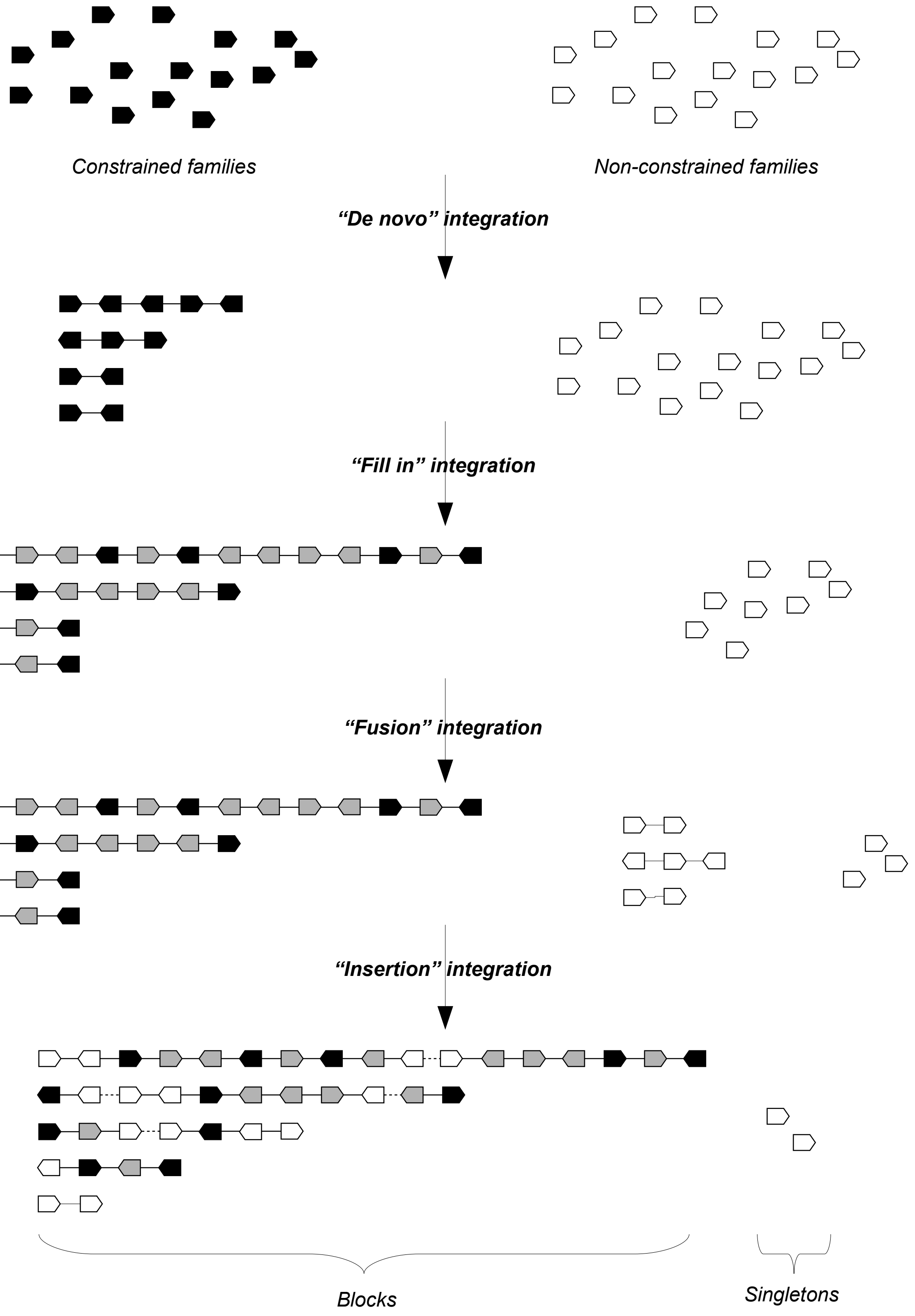

490 **Fig. S8**

Multi-integration summary. All four integrations are run in a specific order, starting with "de novo" on the constrained genes in order to define the backbone of the blocks. Each further integration combines ancestral genes into blocks or extends existing blocks, gradually increasing the coverage and the precision of the reconstruction.

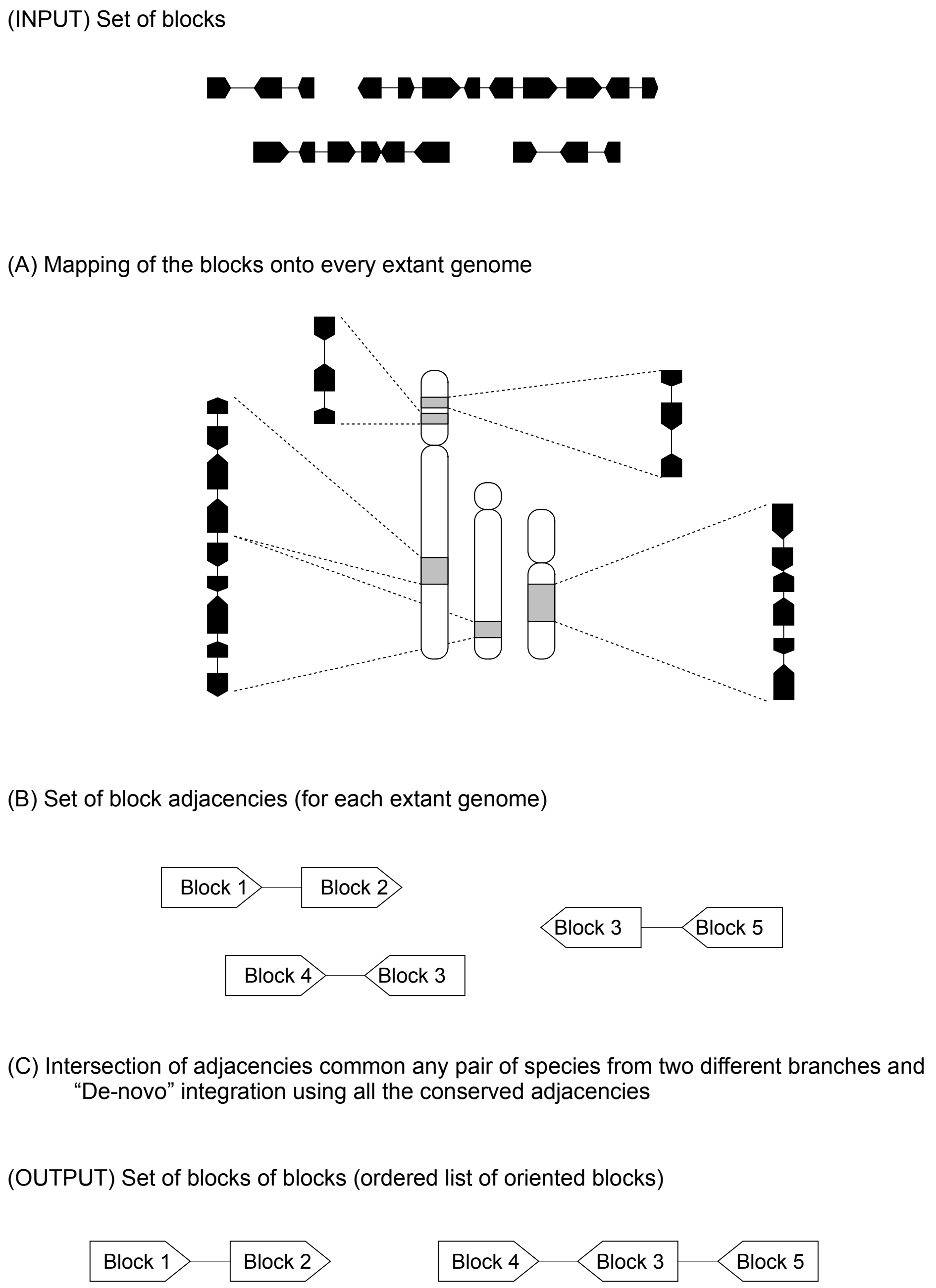

**Fig. S9**

Single-integration on blocks, for a 2-pass reconstruction. Overview of a single-integration reconstruction using blocks instead of genes, thus creating blocks of blocks. While the first pass worked off gene adjacencies, a second pass can be applied by comparing the order of each block in the extant genomes. First, the blocks are mapped onto the extant genomes (possibly on multiple locations), and all the adjacencies of block extremities are extracted. Block adjacencies that are observed across two genomes that cross a given ancestor are added to its weighted adjacency graph, on which the "de novo" integration is run. This second pass is used on all Vertebrates reconstructions (red path on Fig. S1).

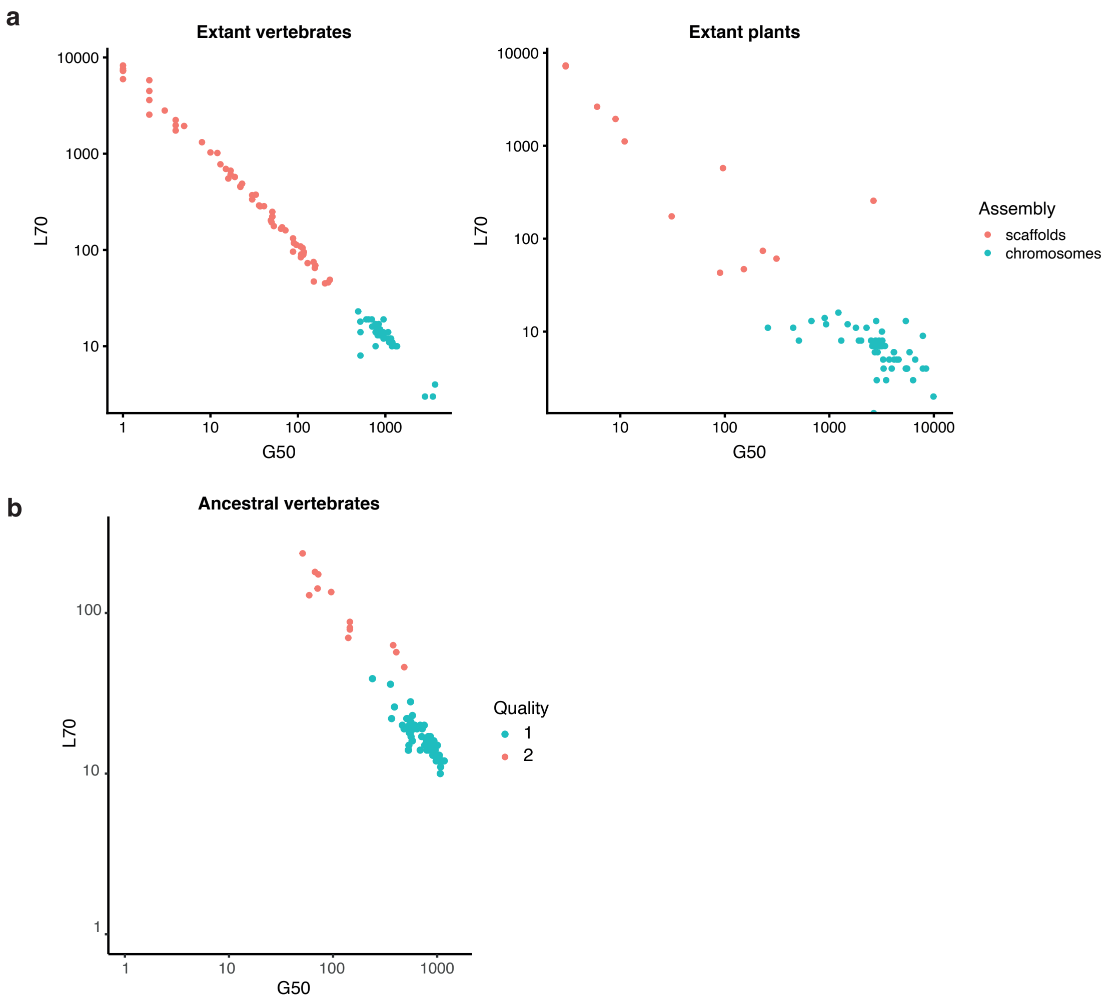

**Fig. S10**

<u>Quality of extant and ancestral genomes</u>. **a.** Chromosomal and low-contiguity assemblies (scaffolds) amongst sequenced extant species are readily distinguished based on the G50 and L70 quality metrics explained in Methods. **b**. Similar distribution for ancestral genomes reconstructed by AGORA based on extant genomes from Ensembl version 102. The combination of thresholds L70 < 40 and G50 > 230 distinguishes low (red) from high (blue) quality reconstructions used in analyses.

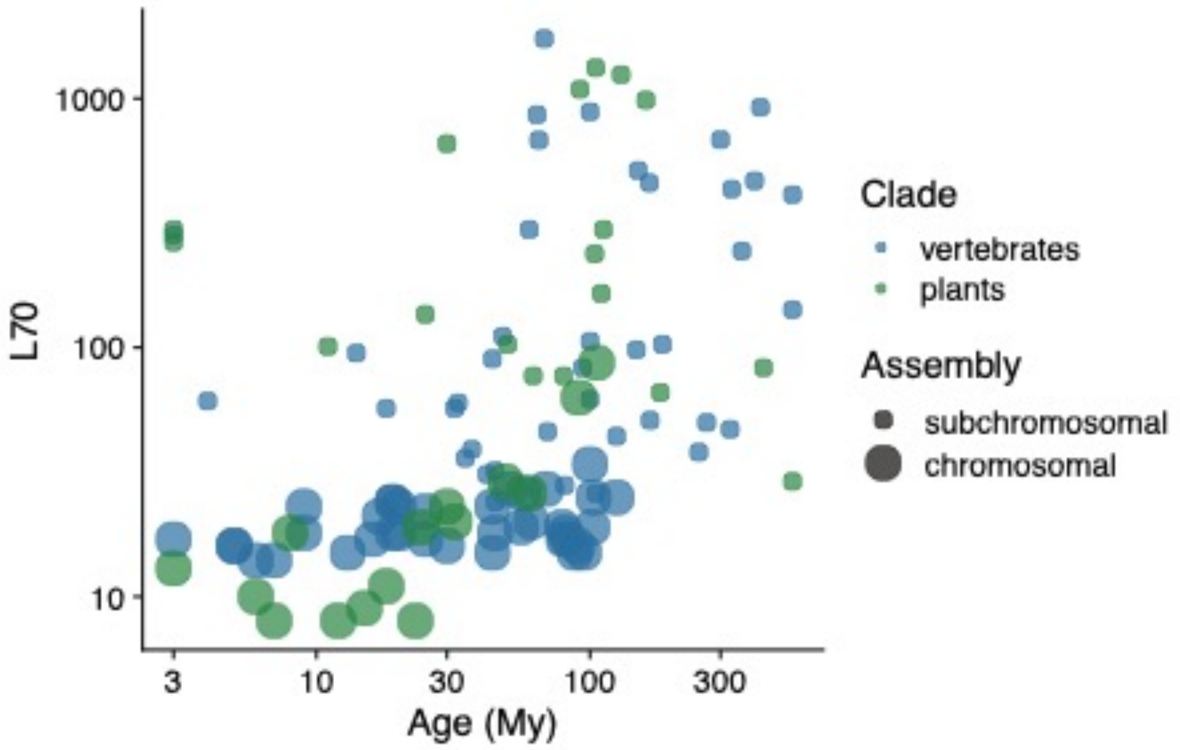

**Fig. S11**

Ancestral genome quality as a function of Age (My). Ancestral genome quality is measured as the L70: as in Fig. S10, the smallest number of blocks adding up to 70% of the total genome length, measured in gene units. Younger ancestors tend to be reconstructed better (longer blocks) than older ones, and can be considered high-quality (chromosome-level) as per the thresholds explained in Fig. S10.

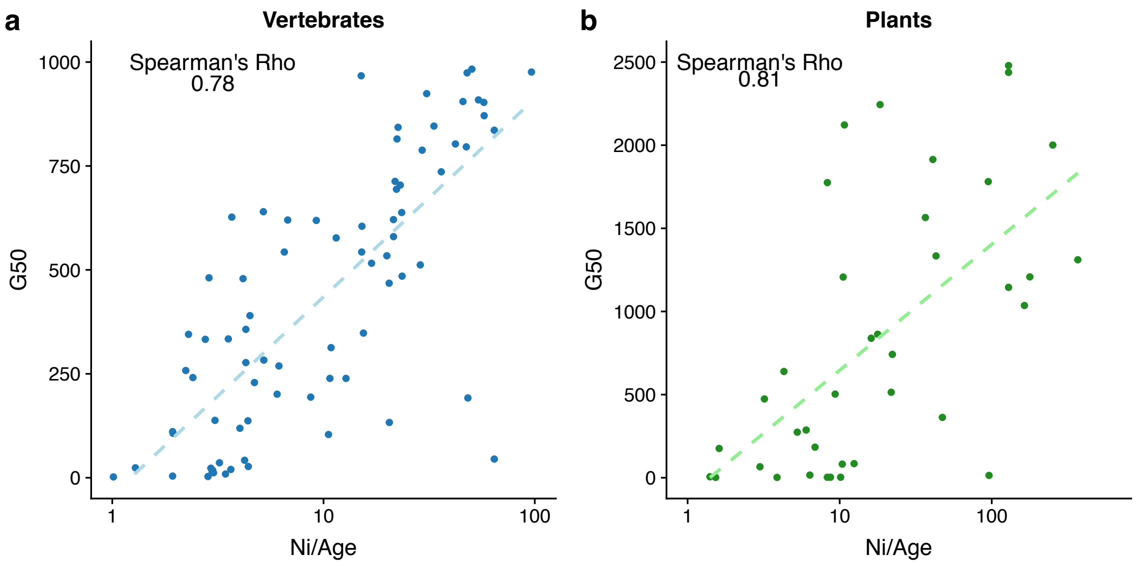

**Fig. S12**

Assembly quality as a function of Ni/Age. Ancestral genome contiguity is related to the number of sequenced extant species informative for this ancestor, and ancestor's age. Correlation of ancestral genome contiguity (G50 metric, see Methods) with the number of informative pairwise comparisons between extant genomes (Ni) normalized by ancestor age (Age, in My), for: **A**. 59 vertebrate ancestral genomes, and **B**. 48 plant ancestral genomes.

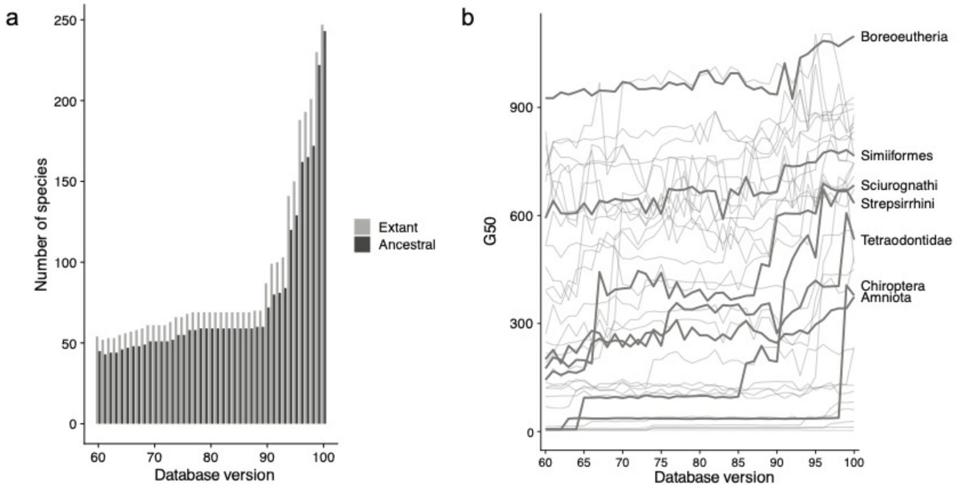

**Fig. S13**

Number of genomes and quality of the ancestral reconstructions in the Genomicus database. The quality of the AGORA reconstructions, measured with the G50 metric (see Methods), have overall increased throughout the versions of Genomicus, as a result of new extant genomes being added to the Ensembl database. Since the version 90 of Genomicus (based on Ensembl 90), the number of genomes has been steadily rising at a fast pace, and is causing a faster improvement of ancestral genomes quality.

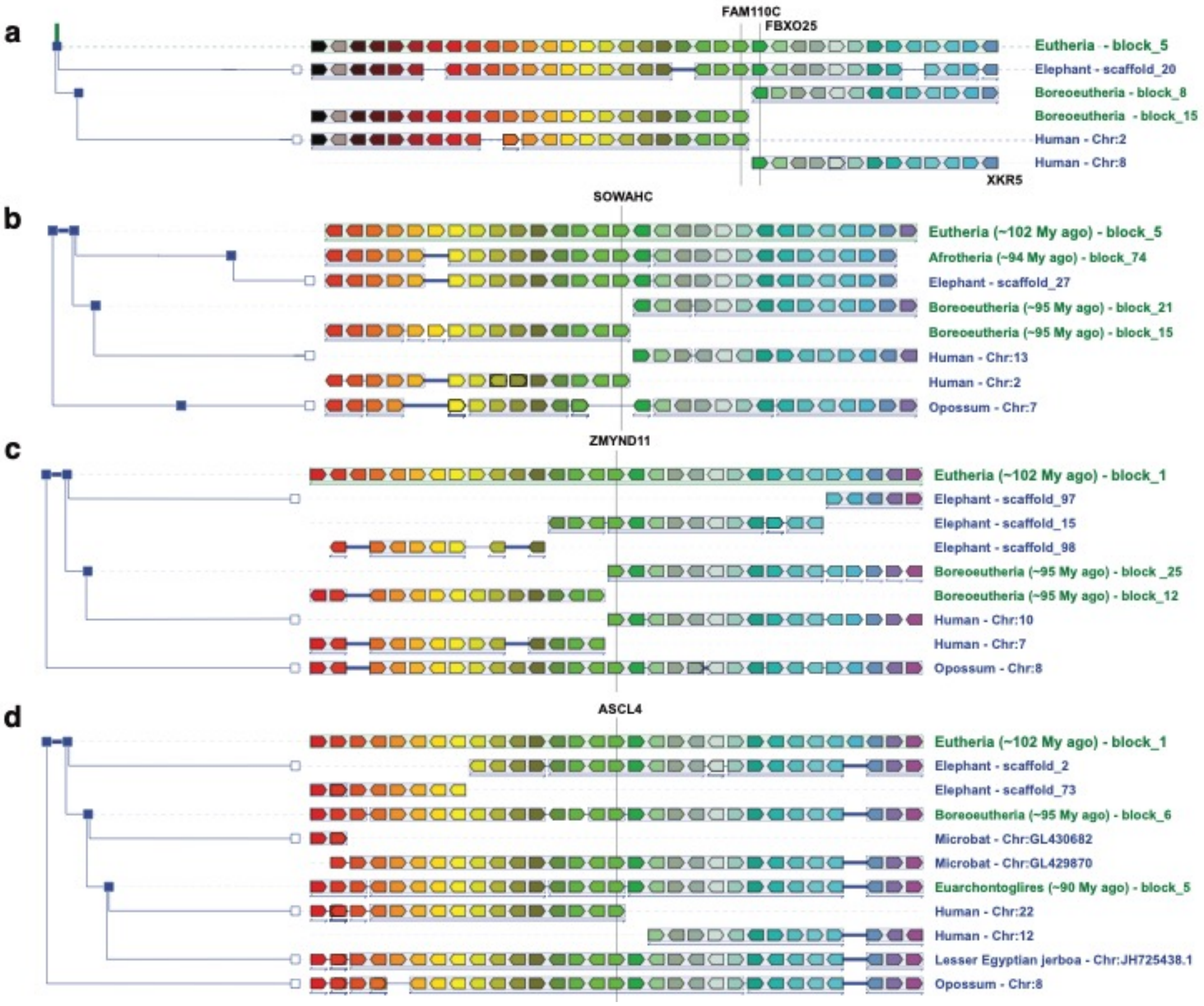

**Sup. Fig. 14**

Simplified views of Genomicus v92 AlignView show gene adjacencies in extant species that support the ancestral linkage of human chromosomes 8-2-13, 7-10, 12-22 in *Eutheria*. Alignments between gene orders in ancestral (*Eutheria*, *Boreoeutheria*) and extant species (human, elephant, opossum…). Orthologs or ancestors of each extant gene are shown in matching colours. **A.** *Eutheria* CAR-5 corresponds to an ancestral linkage between human chromosomes 8 and 2, and the ancestral adjacency of genes FAM110C and FBXO25 is conserved in elephant. **B**. *Eutheria* CAR-5 also includes an ancestral linkage of human chromosomes 13 and 2, and the ancestral configuration is supported by the conserved gene neighbourhood in elephant and opossum. **C**. *Eutheria* CAR-1, an ancestral linkage between human chromosomes 7 and 10, is supported by homologous neighbors genes of ZMYND11 in elephant and opossum. **D**. *Eutheria* CAR-1, *Boreoeutheria* CAR-6 and *Euarchontoglires* CAR-5 correspond to an ancestral linkage between human chromosomes 22 and 12, supported by the neighbour genes of ASCL4 in elephant, microbat, jerboa and opossum.

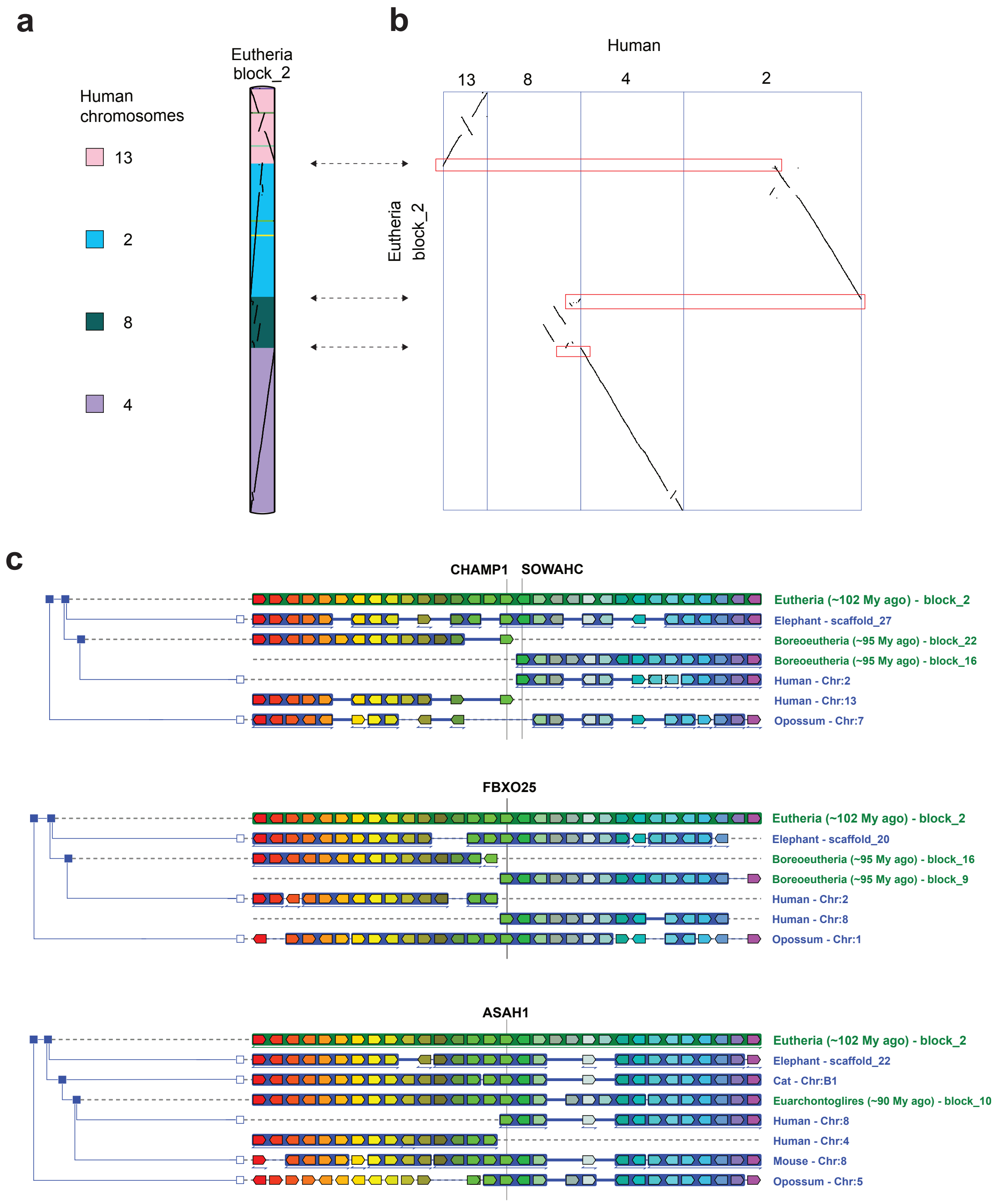

**Fig. S15**

Evidence for ancestral linkage of human chromosomes 4, 8, 2 and 13 in *Eutheria*. Our reconstruction of the *Eutheria* ancestral genome with Ensembl v.92 data resulted in two ancestral CARs corresponding to segments of human chromosomes 4, 8, 2, and 13: chr. 4 and 8 in one, and chr. 8, 2, and 13 in the other. Our later reconstruction using Ensembl v.102 data now links both segments and infers a single ancestral CAR (block_2), like DESCHRAMBLER, as shown in the chromosome painting view (a), the gene order comparison (b), and the Genomicus AlignView (c). All three junctions are supported by the elephant (ingroup) and the opossum (outgroup), though on different chromosomes or scaffolds each time. Some ancestral junctions, such as between human chromosomes 4 and 8, are supported by other species too.

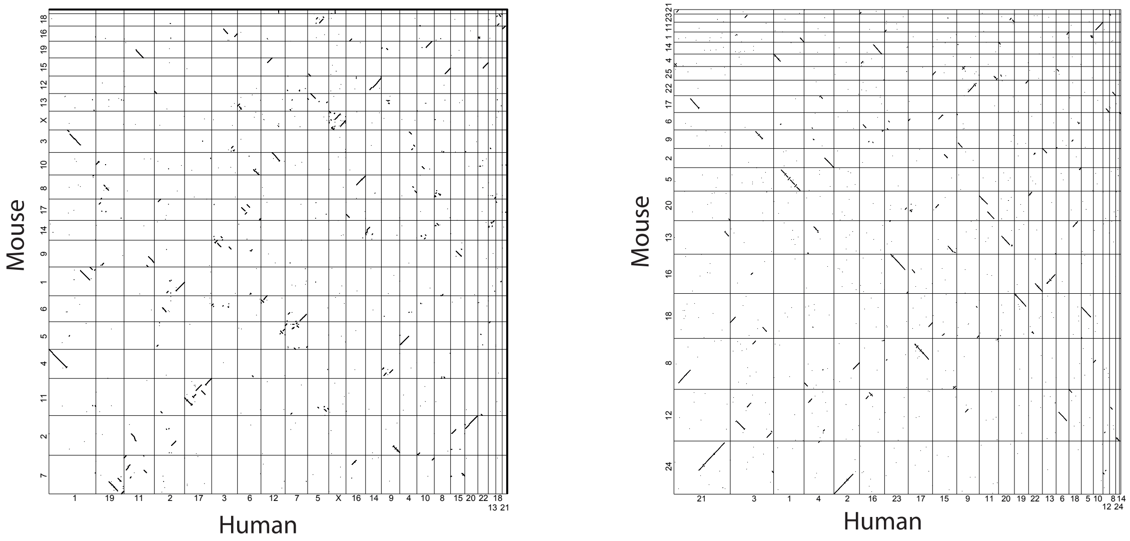

Real genomes

Simulated genomes

**Fig. S16** Genome comparisons in real and simulated genomes. The left panel shows a dot matrix of human (x-axis) versus mouse (y-axis) homologous genes from Ensembl version 102. Numbers on the axes indicate chromosomes in the respective genomes, and genes coordinates are their rank in each chromosome. The right panel shows a comparison between one each of the 50 simulated human and mouse genomes that were used to benchmark AGORA. The comparison shows that the complexity of the simulation is high, breaking chromosomes and dispersing genes in a way that is similar to the real evolution of the human and mouse genomes.

### References

1. Cunningham, F. *et al.* Ensembl 2022. *Nucleic Acids Res* gkab1049 (2021) doi:10.1093/nar/gkab1049.
2. https://www.ensembl.org/info/genome/compara/homology_types.html#paralogues.
3. Kim, J. *et al.* Reconstruction and evolutionary history of eutherian chromosomes. *Proc. Natl. Acad. Sci. U.S.A.* **114**, E5379–E5388 (2017).
4. Lucas, J. M. & Roest Crollius, H. High precision detection of conserved segments from synteny blocks. *PLoS ONE* **12**, e0180198 (2017).
5. Kumar, S., Stecher, G., Suleski, M. & Hedges, S. B. TimeTree: A Resource for Timelines, Timetrees, and Divergence Times. *Molecular Biology and Evolution* **34**, 1812–1819 (2017).
